# Protein Language Models: Is Scaling Necessary?

**DOI:** 10.1101/2024.09.23.614603

**Authors:** Quentin Fournier, Robert M. Vernon, Almer van der Sloot, Benjamin Schulz, Sarath Chandar, Christopher James Langmead

## Abstract

Public protein sequence databases contain samples from the fitness landscape explored by nature. Protein language models (pLMs) pre-trained on these sequences aim to capture this landscape for tasks like property prediction and protein design. Following the same trend as in natural language processing, pLMs have continuously been scaled up. However, the premise that scale leads to better performance assumes that source databases provide an accurate representation of the underlying fitness landscape, which is likely false. By developing an efficient codebase, designing a modern architecture, and addressing data quality concerns such as sample bias, we introduce AMPLIFY, a best-in-class pLM that is orders of magnitude less expensive to train and deploy than previous models. Furthermore, to support the scientific community and democratize the training of pLMs, we have open-sourced AMPLIFY’s pre-training codebase, data, and model checkpoints.

## Introduction

Understanding the relationship between protein sequence and function is a fundamental goal in biology [Dimitrov, 2012, Arnold, 2018]. With the rise of structural biology, this pursuit has often been intertwined with the protein folding problem [Anfinsen, 1973], approaching the challenge as a linkage between sequence-to-structure and structure-to-function relationships. Early progress in this field largely relied on physical modeling based on atomic coordinates, operating under the assumption that predicting enthalpy across specific states would be sufficient to capture the energy landscape of protein folding. However, as the predictive accuracy of these methods began to plateau [Moult et al., 2011, 2014], it became evident that this assumption was flawed [Wang et al., 2011], highlighting the need for an understanding of configurational entropy. This realization has driven the search for a new mathematical framework to study protein function.

Recent breakthroughs in using machine learning approaches to address protein structure prediction and design have emerged as a direct consequence of this shift [Senior et al., 2020, Jumper et al., 2021, Dauparas et al., 2022, Wang et al., 2022, Ingraham et al., 2023, Watson et al., 2023, Abramson et al., 2024]. The field of information science has a long-standing tradition of handling entropy as a discrete measurement of probability states, and the mathematical foundations of modern machine learning (ML) are naturally suited to model the dynamic, multi-state behavior of proteins [Kamisetty et al., 2011]. However, many studies continue to be constrained by a structure-based definition of protein function, with models primarily focused on the subset of sequence space represented in databases of experimentally determined atomic coordinates.

While understanding structure-to-function relationships remains crucial, recent discoveries in molecular biology have shown that protein sequence influences function far beyond just protein folding. The dynamic nature of proteins, especially intrinsically disordered proteins, is key to both cellular and extracellular organization. Protein sequences have significant macroscale and observable impacts on the formation of organelles, materials, and cellular systems governed by complex phenomena such as solvent interactions, dynamic states, transient populations, and the fluid assembly of proteins as polymeric chains and colloids [Mittag et al., 2010, Muiznieks and Keeley, 2013, Banani et al., 2017, Shin and Brangwynne, 2017, Sabari et al., 2018, Shin et al., 2018, Quiroz et al., 2020, Lafontaine et al., 2021, Xiao et al., 2022].

Sequence databases avoid the biased sampling inherent to structure databases, and by approaching protein sequence as a natural language problem, researchers have demonstrated that protein language models (pLMs) can learn structural information independently of structure databases [Rives et al., 2021]. These models capture structural information through statistical patterns that emerge when protein folding acts as part of the physical ‘grammar’ guiding the evolution of protein sequences [Eddy, 1998, Vig et al., 2020].

Here, we focus on Transformer-based pLMs [Rives et al., 2021, Lin et al., 2023], which are trained solely on protein sequences and are frequently used in the context of property prediction and generative design [Wittmann et al., 2021, Ferruz and Höcker, 2022].

Since their introduction, the trend has been to scale the size of pLMs by increasing the width and depth of the neural networks. Modern sequence-only foundation pLMs such as ESM2 and ProGen2 perform best at 15 billion and 6.4 billion parameters, respectively. More recent multimodal models, such as ESM3, have been scaled even more to a staggering 98 billion parameters. This scaling is motivated by the recent achievements of larger networks in natural language processing, where their language modeling abilities have been shown to scale as a power law of the model size given a dataset and compute budget [Kaplan et al., 2020, Hoffmann et al., 2022]. However, it is worth asking: *Is scaling necessary to improve pLM performance?* A negative answer to that question could democratize pLM development, as existing foundation models are prohibitively expensive to pre-train for most research labs. For example, training ESM2 15B required an estimated 8.1×10^22^ FLOPs, costing approximately 1.5 million USD on AWS, excluding the cost of training intermediate models required for its development [Hoffmann et al., 2022]. The cost of pre-training pLMs means that, for most scientists, the only way to contribute to the field is through fine-tuning and other forms of model alignment. Finally, it is important to note that large models are not only costly to train but also to use, limiting the computational efficiency of future work built on them.

There are three strategies to enhance a pLM’s performance without altering its architecture or size: (i) increase data quality, (ii) increase data quantity, and (iii) increase the number of training steps. We evaluated all three strategies while incorporating recent algorithmic advances to reduce the computational requirements of training. Our findings show that all three strategies are effective, but increasing data quality alone can yield state-of-the-art pLMs at a much smaller scale than existing foundation models. In particular, our best model, AMPLIFY 350M (**A**mgen-**M**ila **P**rotein **L**anguage model for **I**n**F**erence and discover**Y**), outperforms the best sequence-only pLM, ESM2 15B, yet has 43× fewer parameters, requires 17× fewer FLOPs to train, and achieves 24 to 29× higher throughput at inference time, depending on sequence length. This aligns with findings in other domains, where careful curation of training data often competes well with scale [Gao et al., 2020, Paul et al., 2021, Sorscher et al., 2022, Penedo et al., 2023, 2024].

We compare the performance of this sequence-only model to structure-based models on specific tasks, demonstrating that models trained on structural data do not perform well across the full distribution of natural proteins. In our predictions, we find that AlphaFold2 [Bryant et al., 2022] struggles to distinguish between non-proteins and disordered proteins, whereas AMPLIFY exhibits an emergent ability to do so with high accuracy. Sequence-to-structure and structure-to-sequence models are commonly used tools in the field [Dauparas et al., 2022], but our benchmarking shows that they do not match the performance of pLMs on two critical tasks relevant to protein design.

To promote the democratization of pLM development, we make available our training and test sets, the script to reproduce them from public sources, the training code, and the AMPLIFY models. All of these resources are released under the permissive open-source MIT license.

## Results

We sought to compare the effects of adjusting pLM size against other controllable factors: (i) the contents of the training and test datasets, (ii) their sizes, and (iii) the number of training steps. As detailed in the Methods section, our choices for (i) differ from prior work primarily in three ways: (a) our training set is derived from UniRef100 and supplemented with paired antibody sequences from the Observed Antibody Space (OAS) [Kovaltsuk et al., 2018] and individual domains from the Structural Classification of Proteins (SCOP) database [Chandonia et al., 2022], whereas prior work uses UniRef50/90 sequences; (b) we use protein evidence level annotations (UniProt 2023) to select high confidence candidates for our test set; and (c) proteome completeness scores [Simão et al., 2015, Seppey et al., 2019] to ensure that our test set is selected by a random split out of a natural distribution (i.e., sequences are selected proportional to their abundance in full proteomes). These changes aim to make the test set more representative of the fitness landscape by including only high-confidence sequences in their natural distribution as found in complete proteomes.

We based our AMPLIFY model on the architecture of the previous state-of-the-art sequence-only LLM, ESM2 [Lin et al., 2022], and implemented separate data pipelines to replicate prior workflows and our own choices for factors (i) and (ii) above. In total, 21 models were trained and evaluated, amounting to 1.68×10^22^ FLOPs of compute, and an additional 18 pre-trained models from public sources were evaluated against our test sets.

### Data Curation Improves Protein Likelihood and Recovery Independent of Model Size and Training Duration

To measure the impact of model size and of training set composition, we pre-trained 21 models from scratch: ESM2 8M, 35M, and 150M trained on UniRef50, AMPLIFY 120M on UniRef50, UniRef100, as well as every consecutive yearly release of UniRef100 since 2011, and AMPLIFY 120M and 350M on our dataset. Reproducing the larger ESM2 is prohibitively costly as a single ESM2 15B would require more compute than all the aforementioned models. Pre-trained versions of ESM2 and ProGen2 [Nijkamp et al., 2023] models were also obtained from public sources to serve as controls and to confirm that our pre-training code produced equivalent models. Performance is measured in terms of pseudo-perplexity and sequence recovery accuracy. Pseudo-perplexity measures the likelihood of data under a model, that is, a model with lower perplexity assigns a higher likelihood to the proteins in the dataset. Assuming the data reflects the distribution of natural proteins, which our test set aims to, then a model with lower perplexity has a greater confidence in identifying natural proteins. Sequence recovery evaluates whether the residue type predicted by the model for a masked position matches the actual (or consensus) residue, and a higher accuracy at this task is useful for protein engineering. In all cases, these metrics are reported on our held out test set. However, we cannot exclude the possibility that the training set of the public models may have included some of our test sequences, leading to their performance metrics being artificially inflated. Figure 1A shows that larger models learn faster than smaller ones, but also that models trained on UniRef100 plus OAS antibody sequences and SCOP domains (UR100P^1^) learn faster than those on UniRef50, likely due to UR100P’s greater sequence quantity and diversity.

**Figure 1:**
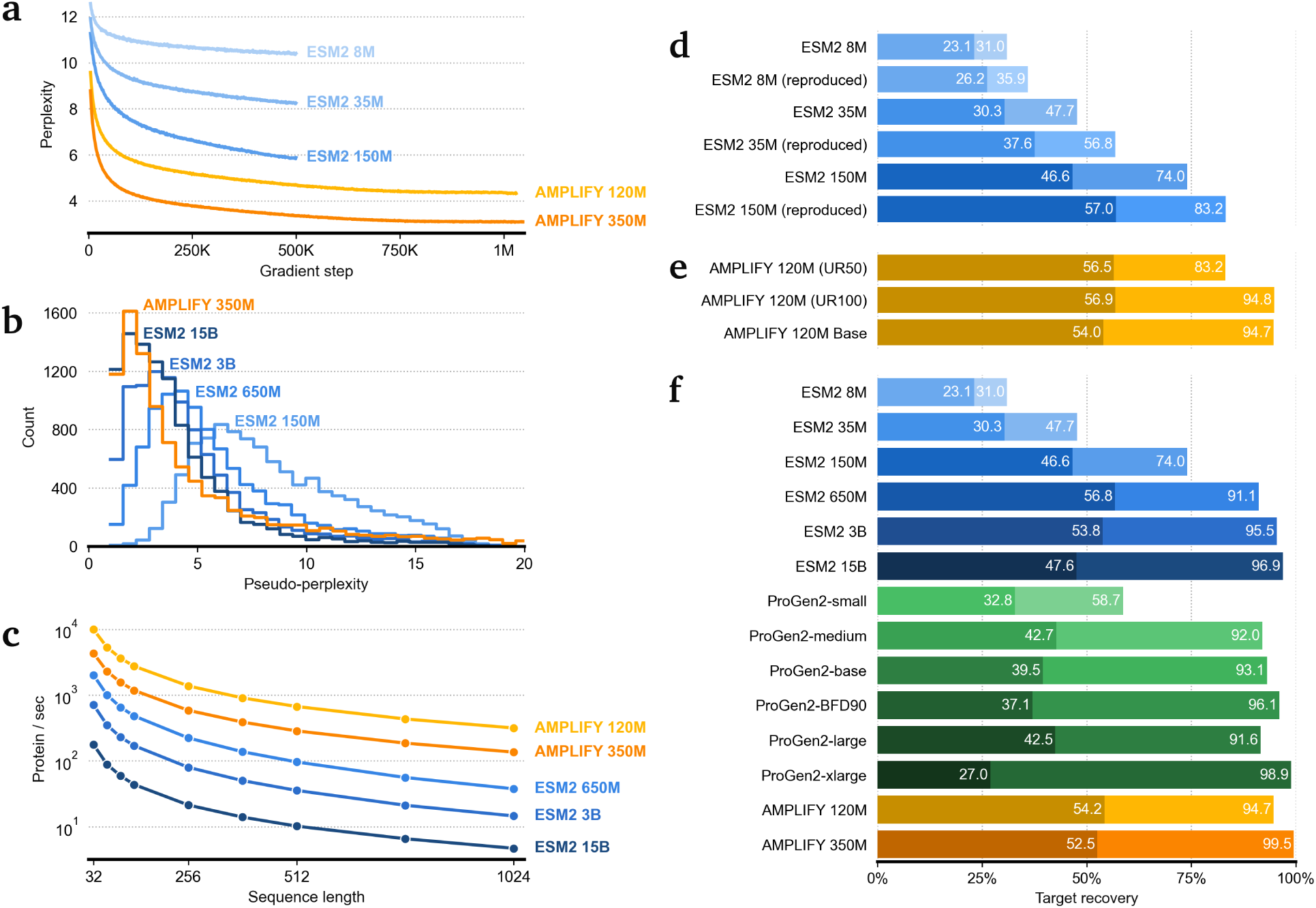
Data curation improves convergence speed, the likelihood of natural proteins under the model, as well as sequence recovery, independent of model scale or training duration. A) Test perplexity on UniProt for ESM2 controls in blue (8M, 35M, and 150M, from light to dark) and AMPLIFY in orange (120M and 350M, from light to dark), demonstrating that model convergence does not occur at 500K steps, even for smaller models. B) Test pseudo-perplexity on UniProt for ESM2 in blue (150M, 650M, 3B, and 15B, from light to dark) and AMPLIFY 350M (orange), with the latter showing comparable performance to ESM2 15B despite a 43× reduction in scale. C) Inference efficiency measured as proteins per second as a function of sequence length for ESM2 in blue (150M, 650M, 3B, and 15B, from light to dark) and AMPLIFY in orange (120M and 350M, from light to dark). D-F) Sequence recovery accuracy on human proteome binned by sequence conservation. Light bars show recovery of the target sequence at positions with 96 to 99% sequence conservation and shaded bars show recovery of the most common non-target sequence when found at >70% conservation. The gap between them highlights the degree of undesirable memorization.

Figure 1B shows the non-smoothed test pseudo-perplexity histograms for a range of pre-trained ESM2 and our best model, AMPLIFY 350M. As expected, both the position and width of the modes decrease as the size of the model increases. However, it is noteworthy that a 350M parameter AMPLIFY model is competitive with, and arguably better than, a 15B parameter ESM2 model at modeling the distribution of unseen natural proteins.

Figures 1D-F show similar trends on a different test set, the human proteome, and performance metric, the sequence recovery accuracy at positions with high sequence conservation. As before, performance increases with the size of the model, but the contents of the training and test sets also have large effects on performance. Taken together, we conclude that data curation can significantly improve model performance, independent of model size or training duration.

### Clustering Sequences to Reduce Similarity in Training Data Harms Performance

Contrary to Lin et al. [2023], we found that perplexity continued to decrease beyond 500K updates, up to at least 1M training steps (Figure 1A). This difference can be attributed to the differences in the data curation decisions made during the construction of training and test sets. While Lin et al. [2023] used a subset of UniRef50 for test regardless of whether UniRef50 or UniRef90 was used for training, our training and test sets are derived from UniRef100. The key difference between UniRef50, UniRef90, and UniRef100 is the threshold of allowed sequence similarity. The key difference between UniRef50, UniRef90, and UniRef100 is the threshold of allowed sequence similarity, which are obtained through clustering and with UniRef100 allowing for the greatest amount of similarity. If we assume that most sequences in the various UniRef sets are real in the sense that they are not pseudo-genes or the result of sequencing errors, then UniRef100 is a more representative sample of the fitness landscape. This is because redundancy is a sign of observation count, which we should treat as a measure of confidence that the cluster represents a real part of sequence space. Clustering is in fact expected because of the nature of protein evolution – these clusters represent families of related proteins and they are common for functional reasons. Removing sequences that show up in large clusters downweights confident sequences and ultimately upweights singleton sequences that are more likely to derive from sequencing errors and other sources of noise. While we acknowledge that singletons could be rare yet functional proteins, the large number of entries removed every year from the databases suggests most of them might be non-proteins (Figure 2F). We thus hypothesize that the saturation observed by Lin et al. [2023] at 500K steps is due to their use of a UniRef50-based test set, which lacks the natural sequence diversity required to identify that reducing redundancy can harm model performance. Further evidence in support of the hypothesis that UniRef100 represents a more natural sequence distribution can be observed by the increase of ambiguous sequence reads, fragmentary sequences, and hypothetical proteins in UniRef50 compared to UniRef100, as well as the significant depletion of specific functional classes (Table 5).

**Figure 2:**
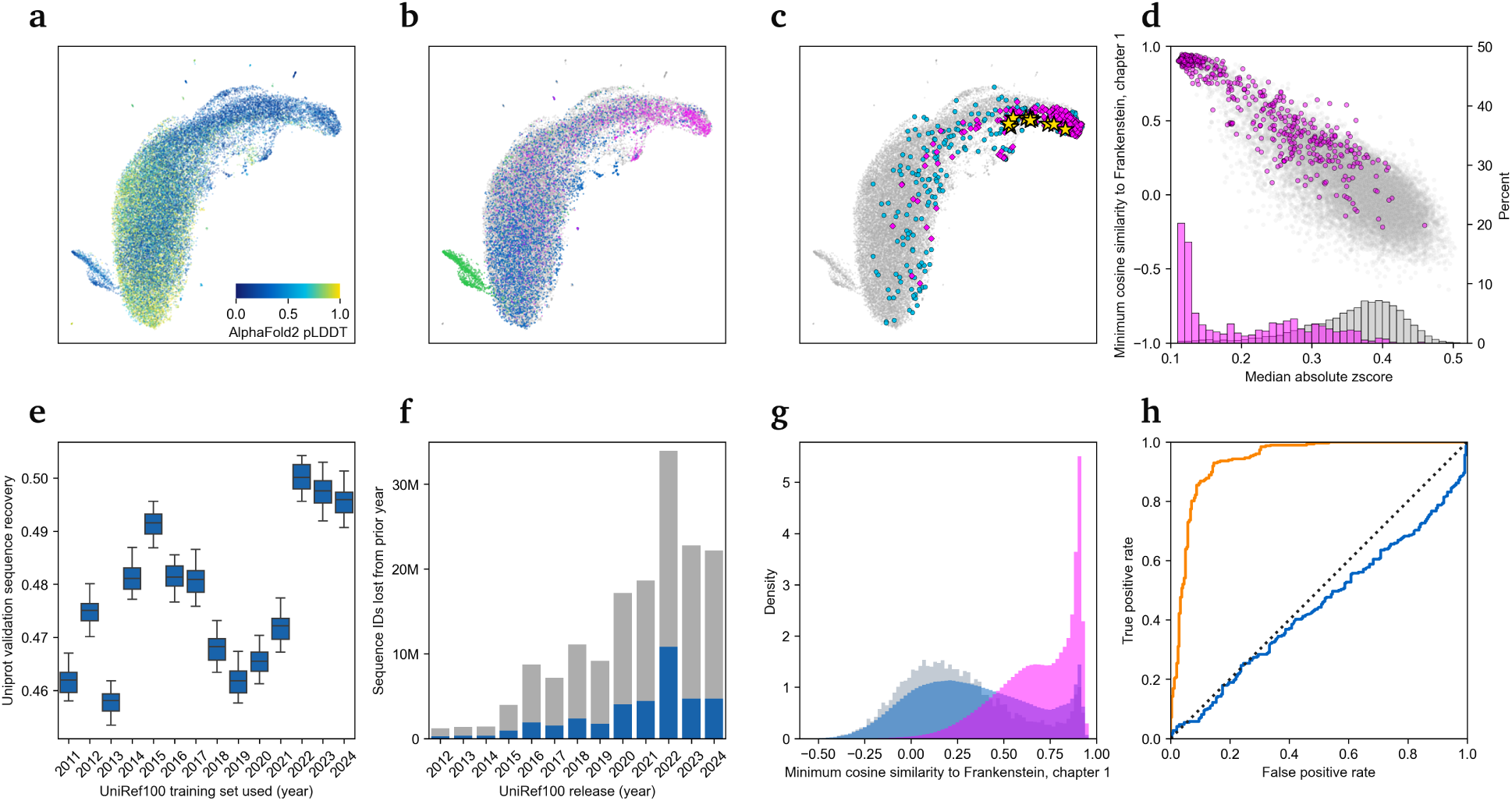
Emergent properties. A-C) RG motifs of the human proteome embedded with AMPLIFY 350M and projected with UMAP. A) Motifs are colored according to their AlphaFold2 pLDDT, highlighting that positions with low structure confidence are widely spread but cluster in two prominent locations. B) Motifs are colored according to three gene ontology term labels: collagen trimer in green, RNA binding in blue, and proteins that do not have any localization annotation in magenta. C) Hypothetical proteins (PE=5) with and without localization annotations are in blue and magenta, respectively. RG motifs in the text of Mary Shelley’s Frankenstein are in yellow and colocalize to the same space as low evidence, unannotated proteins. D) The minimum per-residue similarity of human proteins to Frankenstein against the degree to which residue embeddings diverge from the mean embedding for each full sequence. Proteins with PE=1 and PE=5 are in gray and magenta, respectively. The upper-left cluster indicates that many hypothetical proteins converge to the same space as non-proteins across all residues. E) Sequence recovery on UniProt for AMPLIFY 120M trained on UniRef releases spanning 2011-2024, ranges show performance during the last fifty thousand training steps. A large jump in performance is observed between 2021 and 2022. F) Count of sequence IDs removed between consecutive yearly UniRef releases, with the total count in gray and the count of those with a similarity to Frankenstein >70% in blue. An unusually large cull of sequences happened in 2022, with a high proportion of these at high risk of being non-proteins. G) The distribution of similarity to Frankenstein of our UniProt-based test set in gray matches that of UniRef100 in blue, while the distribution of UniRef50 in magenta is biased towards potential non-proteins. H) The ROC curve of discriminating human proteins annotated as being >25% intrinsically disordered against human proteins with PE=5 and no annotation for localization of AMPLIFY 350M in orange (similarity to Frankenstein) and AlphaFold2 in blue (confidence). AlphaFold2 confidence is clearly incapable of discriminating between disordered proteins and likely non-proteins.

**Table 1:** The full configuration details of AMPLIFY.

|  | AMPLIFY 120M | AMPLIFY 350M |
| --- | --- | --- |
| hidden-size | 640 | 960 |
| num-hidden-layers | 24 | 32 |
| num-attention-heads | 10 | 15 |
| intermediate-size | 2560 | 3840 |
| max-position-embeddings | 2048 | 2048 |
| vocab-size | 27 | 27 |
| rope-theta | 10,000 | 10,000 |
| dropout-prob | 0 | 0 |
| embedding-init-range | 0.02 | 0.02 |
| norm-eps | 1.0e-05 | 1.0e-05 |
| hidden-act | swiglu | swiglu |
| pre-activation-layer-norm | true | true |
| layer-norm-after-embedding | false | false |
| layer-norm-before-last-layer | true | true |
| rms-norm | true | true |
| ffn-bias | false | false |
| attn-bias | false | false |

**Table 2:** Pre-training dataset of each model.

| Model | Source | Major Processing Step | Size | Collection Date |
| --- | --- | --- | --- | --- |
| ESM2 (Original)<br>(8M, 35M, 150M, 650M, 3B, 15B) | UniRef50 | Drawn from UniRef90 | 60M | April 2021 |
| ESM2 (Reproduced)<br>(8M, 35M, 150M)<br>AMPLIFY (UR50)<br>(120M) | UniRef50 | Filter ambiguous and rare amino-acids | 62M | December 2023 |
| ProGen2 (Standard)<br>(small, medium, base, large, xlarge) | UniRef90<br>BFD30 |  | ~170M<br>~57M | Prior to June 2022 |
| ProGen2-BFD90<br>(large) | UniRef90<br>BFD90 | Filter clusters with less than 3 members | ~170M<br>~340M | Prior to June 2022 |
| AMPLIFY (UR100)<br>(120M) | UniRef100 | Filter ambiguous and rare amino-acids | 370M | December 2023 |
| AMPLIFY<br>(120M Base, 120M, 350M) | UniRef100<br>OAS<br>SCOP | Filter ambiguous and rare amino-acids<br>Filter ambiguous and rare amino-acids<br>Filter ambiguous and rare amino-acids | 370M<br>3.6M<br>24k | December 2023 |

**Table 3:**
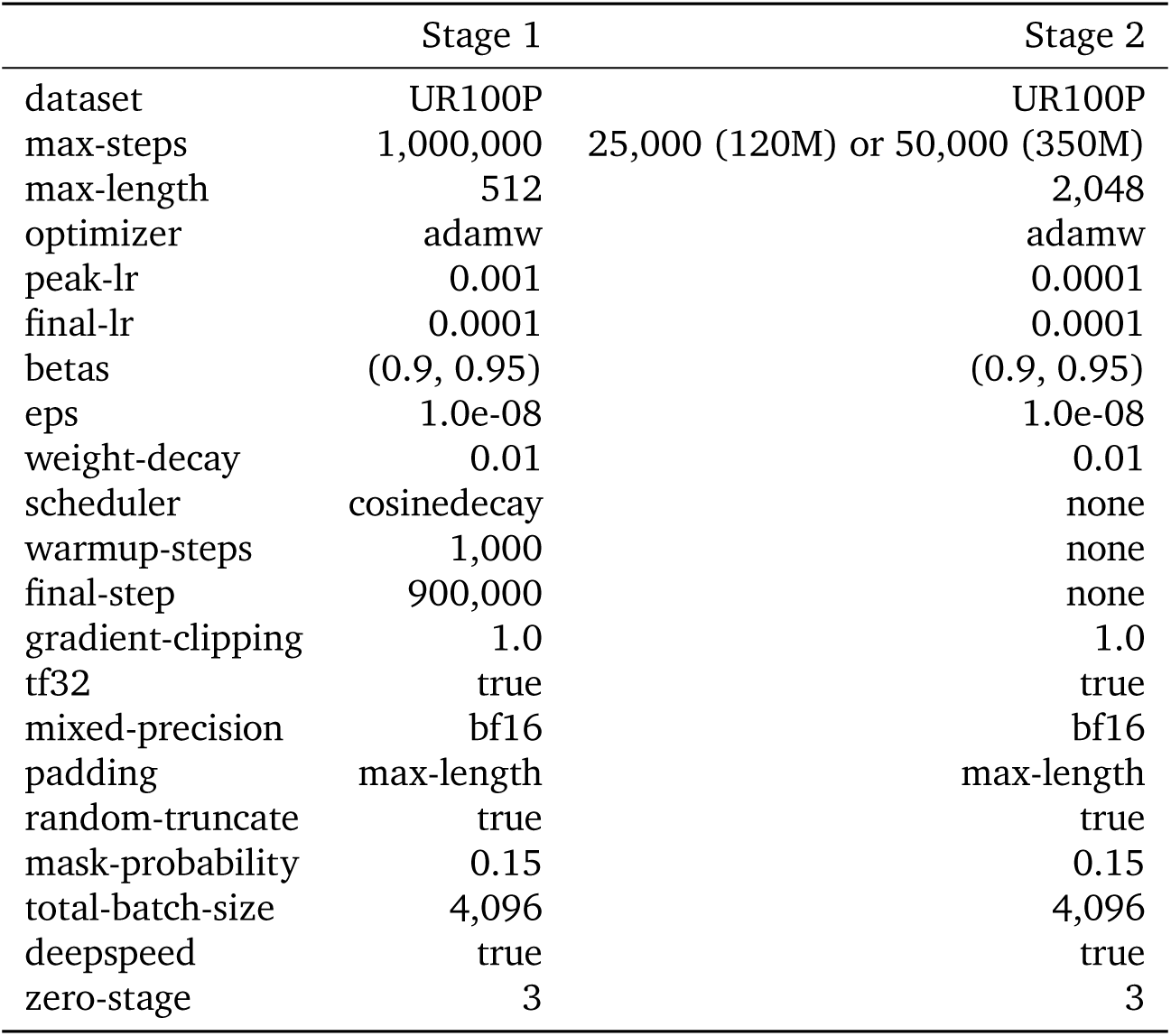
The full configuration details for AMPLIFY’s optimization.

|  | Stage 1 | Stage 2 |
| --- | --- | --- |
| dataset | UR100P | UR100P |
| max-steps | 1,000,000 | 25,000 (120M) or 50,000 (350M) |
| max-length | 512 | 2,048 |
| optimizer | adamw | adamw |
| peak-lr | 0.001 | 0.0001 |
| final-lr | 0.0001 | 0.0001 |
| betas | (0.9, 0.95) | (0.9, 0.95) |
| eps | 1.0e-08 | 1.0e-08 |
| weight-decay | 0.01 | 0.01 |
| scheduler | cosinedecay | none |
| warmup-steps | 1,000 | none |
| final-step | 900,000 | none |
| gradient-clipping | 1.0 | 1.0 |
| tf32 | true | true |
| mixed-precision | bf16 | bf16 |
| padding | max-length | max-length |
| random-truncate | true | true |
| mask-probability | 0.15 | 0.15 |
| total-batch-size | 4,096 | 4,096 |
| deepspeed | true | true |
| zero-stage | 3 | 3 |

**Table 4:**
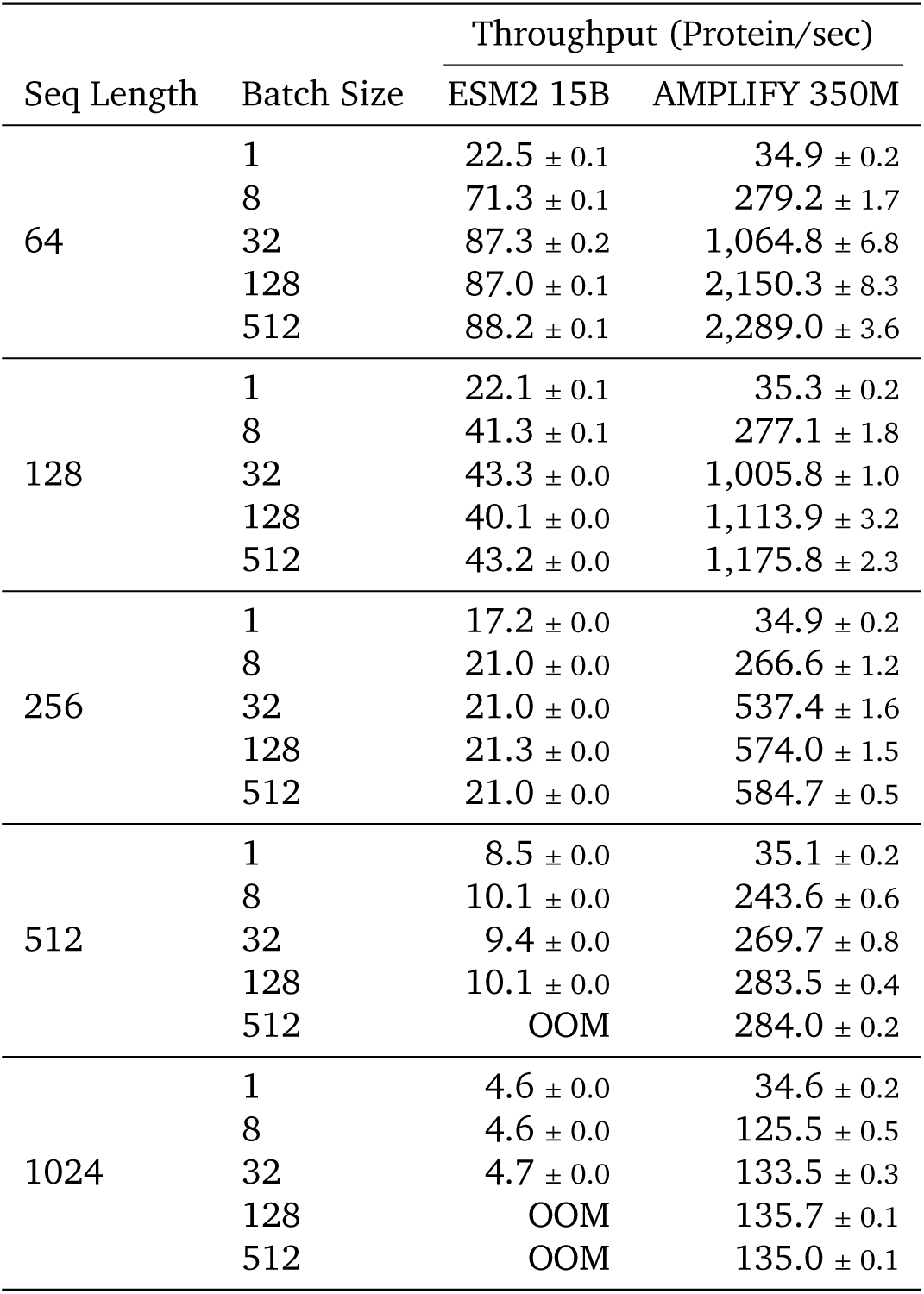
Inference throughput of AMPLIFY 350M and ESM2 15B as measured by the number of proteins per second processed on a single A100 without padding.

| Seq Length | Batch Size | Throughput (Protein/sec) |  |
| --- | --- | --- | --- |
|  |  | ESM2 15B | AMPLIFY 350M |
| 64 | 1 | 22.5 ± 0.1 | 34.9 ± 0.2 |
| | 8 | 71.3 ± 0.1 | 279.2 ± 1.7 |
| | 32 | 87.3 ± 0.2 | 1,064.8 ± 6.8 |
| | 128 | 87.0 ± 0.1 | 2,150.3 ± 8.3 |
| | 512 | 88.2 ± 0.1 | 2,289.0 ± 3.6 |
| 128 | 1 | 22.1 ± 0.1 | 35.3 ± 0.2 |
| | 8 | 41.3 ± 0.1 | 277.1 ± 1.8 |
| | 32 | 43.3 ± 0.0 | 1,005.8 ± 1.0 |
| | 128 | 40.1 ± 0.0 | 1,113.9 ± 3.2 |
| | 512 | 43.2 ± 0.0 | 1,175.8 ± 2.3 |
| 256 | 1 | 17.2 ± 0.0 | 34.9 ± 0.2 |
| | 8 | 21.0 ± 0.0 | 266.6 ± 1.2 |
| | 32 | 21.0 ± 0.0 | 537.4 ± 1.6 |
| | 128 | 21.3 ± 0.0 | 574.0 ± 1.5 |
| | 512 | 21.0 ± 0.0 | 584.7 ± 0.5 |
| 512 | 1 | 8.5 ± 0.0 | 35.1 ± 0.2 |
| | 8 | 10.1 ± 0.0 | 243.6 ± 0.6 |
| | 32 | 9.4 ± 0.0 | 269.7 ± 0.8 |
| | 128 | 10.1 ± 0.0 | 283.5 ± 0.4 |
| | 512 | OOM | 284.0 ± 0.2 |
| 1024 | 1 | 4.6 ± 0.0 | 34.6 ± 0.2 |
| | 8 | 4.6 ± 0.0 | 125.5 ± 0.5 |
| | 32 | 4.7 ± 0.0 | 133.5 ± 0.3 |
| | 128 | OOM | 135.7 ± 0.1 |
| | 512 | OOM | 135.0 ± 0.1 |

**Table 5:** Enrichment analysis of keywords and metadata stored in the protein name field from the source FASTA files.

|  | UniRef100 Count | UniRef50 Count | UniRef50 Enrichment |
| --- | --- | --- | --- |
| UniProt-SPROT | 481,964 | 199,710 |  |
| UniProt-TrEMBL | 203,400,221 | 50,742,156 |  |
| Other | 172,682,262 | 11,818,025 |  |
| Total Size | 376,564,447 | 62,759,891 |  |
| Protein Evidence Level |  |  |  |
| 1. Experimental Evidence At Protein Level | 318,277 | 122,090 | 1.53 |
| 2. Experimental Evidence At Transcript Level | 1,186,593 | 238,615 | 0.80 |
| 3. Protein Inferred From Homology | 72,111,812 | 7,076,681 | 0.39 |
| 4. Protein Predicted | 130,263,740 | 43,502,946 | 1.34 |
| 5. Protein Uncertain | 1,763 | 1,534 | 3.48 |
| Enriched Keywords In Protein Name Fields |  |  |  |
| Uncharacterized | 29,474,903 | 16,775,040 | 2.28 |
| Repeat Protein | 717,606 | 311,832 | 1.74 |
| Intergenic Region | 894 | 351 | 1.57 |
| Fragment | 18,239,506 | 7,136,543 | 1.57 |
| Domain-Containing | 36,560,980 | 14,237,594 | 1.56 |
| Hypothetical Protein | 317,231 | 119,720 | 1.51 |
| Low-Complexity | 6,920 | 2,364 | 1.37 |
| Depleted Keywords In Protein Name Fields |  |  |  |
| Kinase | 5,104,086 | 1,079,096 | 0.85 |
| Polymerase | 1,972,112 | 354,276 | 0.72 |
| Receptor | 1,566,374 | 264,910 | 0.68 |
| Enzyme | 902,609 | 143,035 | 0.63 |
| Transporter | 8,321,056 | 1,018,653 | 0.49 |
| ATP-binding Protein | 1,782,336 | 148,472 | 0.33 |
| Chaperone Protein | 153,827 | 7,712 | 0.20 |

### The Mask Replacement Task May Encourage Overfitting as Models Scale

The primary task for pre-training pLMs is mask replacement, whose objective is to predict the residue type of masked positions chosen at random given the rest of the sequence. One might assume that the theoretically best model would achieve perfect accuracy in recovering these masked positions. However, this is not ideal for two reasons. First, such a model may have overfit the training data by memorizing it. Second, when a pLM is used to design novel sequences, the goal is to generate novel functionally valid designs, which may not always match the residue types in the training or test sets. Thus, a key risk of scaling pLMs is overfitting in the form of undesirable memorization.

To test overfitting in the mask replacement task, we created an orthogonal task using sequence conservation statistics from the human proteome. We categorized individual sequence positions based on their presence in the training data and the amino acid frequencies at matching positions in close homologs (see Methods). This allowed us to assess mask replacement performance according to the functional importance of sequence identity at each position. We focused on two groups of human proteome residues: (i) highly conserved positions where the residue type seen in the training data appears in 96 to 99% of close homologs, indicating likely functional importance, and (ii) positions where the residue type in homologs occurs >70% of the time, but differs from the type observed in the human sequence. The rationale for this second category comes from protein design, where consensus mutations usually improve protein stability without affecting function [Steipe et al., 1994, Lehmann et al., 2000]. That is, there are positions where a residue ‘synonym’ is not only valid but arguably preferred. By identifying positions where the model does not predict the consensus type at these positions, we can test for overtraining in mask replacement.

Figure 1F shows that as models increase in size, they initially improve at recovering both conserved target sequences (type i) and conserved non-target sequences (type ii). However, larger models eventually overtrain on the target sequences, sacrificing performance on the non-target sequences. This suggests that the largest models are exhibiting overtraining in mask replacement, returning target sequences even when a consensus alternative might be preferable. Such memorization has been previously observed in NLPs. Notably, Carlini et al. [2023] reported that within a model family, larger models memorize 2 to 5× more than smaller ones.

We present this as evidence that a 350M parameter pLM is sufficient to saturate the current training task ^2^.

Consequently, to leverage the increased learning capacity of larger models, we may need to improve the data, redefine the task, or develop more effective methods to prevent overfitting.

### pLMs Should Be Retrained Regularly to Leverage Changes in Data Quality and Quantity

We retrained ESM2 models from scratch to ensure our code could produce models comparable to the published versions. Figure 1D shows that our retrained ESM2 models consistently outperform the published versions in the sequence recovery task described earlier. We hypothesized that this improvement is due to training on more recent versions of UniRef50, because UniRef is updated regularly. As the dataset size increases, we expect a corresponding increase in pLM performance. To test this hypothesis, we trained 120M parameter AMPLIFY models for 250K steps on every UniRef100 yearly release since 2011, covering a 33-fold increase in training data, from approximately 12 to 391 million sequences (Table 7).

**Table 6:**
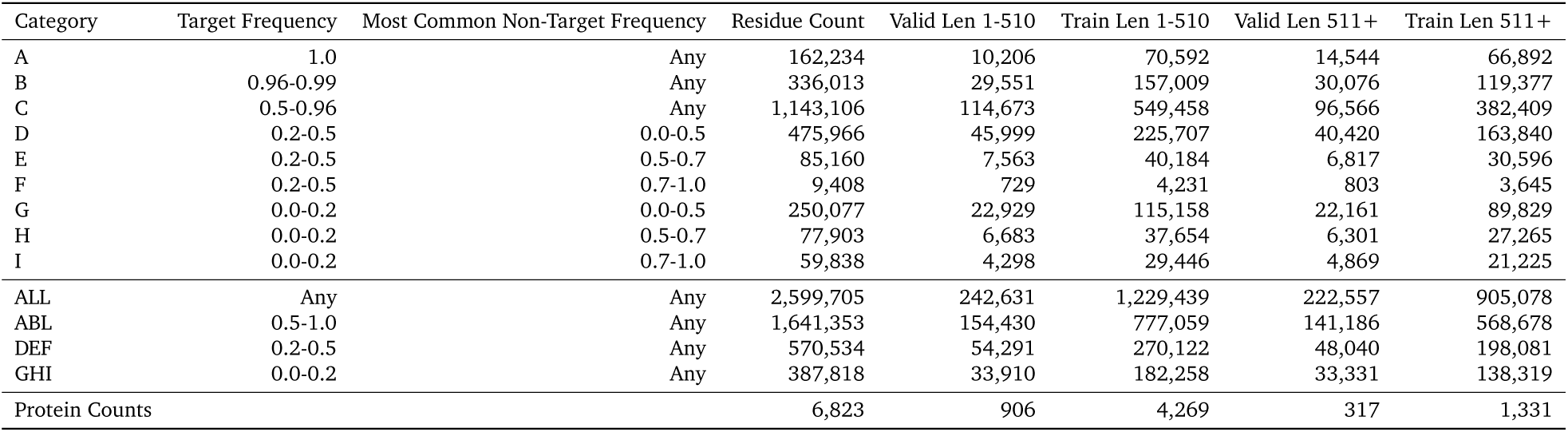
Human proteome test definition.

| Category | Target Frequency | Most Common Non-Target Frequency | Residue Count | Valid Len 1-510 | Train Len 1-510 | Valid Len 511+ | Train Len 511+ |
| --- | --- | --- | --- | --- | --- | --- | --- |
| A | 1.0 | Any | 162,234 | 10,206 | 70,592 | 14,544 | 66,892 |
| B | 0.96-0.99 | Any | 336,013 | 29,551 | 157,009 | 30,076 | 119,377 |
| C | 0.5-0.96 | Any | 1,143,106 | 114,673 | 549,458 | 96,566 | 382,409 |
| D | 0.2-0.5 | 0.0-0.5 | 475,966 | 45,999 | 225,707 | 40,420 | 163,840 |
| E | 0.2-0.5 | 0.5-0.7 | 85,160 | 7,563 | 40,184 | 6,817 | 30,596 |
| F | 0.2-0.5 | 0.7-1.0 | 9,408 | 729 | 4,231 | 803 | 3,645 |
| G | 0.0-0.2 | 0.0-0.5 | 250,077 | 22,929 | 115,158 | 22,161 | 89,829 |
| H | 0.0-0.2 | 0.5-0.7 | 77,903 | 6,683 | 37,654 | 6,301 | 27,265 |
| I | 0.0-0.2 | 0.7-1.0 | 59,838 | 4,298 | 29,446 | 4,869 | 21,225 |
| ALL | Any | Any | 2,599,705 | 242,631 | 1,229,439 | 222,557 | 905,078 |
| ABL | 0.5-1.0 | Any | 1,641,353 | 154,430 | 777,059 | 141,186 | 568,678 |
| DEF | 0.2-0.5 | Any | 570,534 | 54,291 | 270,122 | 48,040 | 198,081 |
| GHI | 0.0-0.2 | Any | 387,818 | 33,910 | 182,258 | 33,331 | 138,319 |
| Protein Counts |  |  | 6,823 | 906 | 4,269 | 317 | 1,331 |

**Table 7:** Uniref100 time course.

| Year | Proteins | Accessions From Uniparc | Accessions From UniProt | Accessions Since Deleted From Uniprot<sup>12</sup>(%) |
| --- | --- | --- | --- | --- |
| 2011 | 11,659,890 | 1,066,364 | 10,593,526 | 22.899 |
| 2012 | 15,688,960 | 1,033,296 | 14,655,664 | 25.593 |
| 2013 | 20,491,135 | 1,139,275 | 19,351,860 | 28.415 |
| 2014 | 33,613,080 | 6,456,416 | 27,156,664 | 30.130 |
| 2015 | 50,371,270 | 14,189,816 | 36,181,454 | 31.144 |
| 2016 | 72,946,704 | 26,119,778 | 46,826,926 | 25.607 |
| 2017 | 94,756,963 | 34,336,823 | 60,420,140 | 23.569 |
| 2018 | 133,853,533 | 43,845,447 | 90,008,086 | 21.415 |
| 2019 | 172,327,164 | 55,571,957 | 116,755,207 | 19.195 |
| 2020 | 216,491,817 | 68,241,824 | 148,249,993 | 16.487 |
| 2021 | 262,115,656 | 87,560,569 | 174,555,087 | 18.838 |
| 2022 | 297,827,854 | 103,262,868 | 194,564,986 | 12.210 |
| 2023 | 342,650,445 | 131,626,082 | 211,024,363 | 8.578 |
| 2024 | 390,790,959 | 174,302,933 | 216,488,026 | 4.224 |

Figure 2E shows that the relationship between UniRef releases and model performance is not straightforward. From this analysis, we obtained three major findings. First, performance on our sequence recovery tasks does not significantly benefit from the additional data added over time (Figure 5). Second, as expected, the main benefit of the additional data is in preventing overtraining. This is most evident in our orthogonal task, where the addition of data enhances the model’s ability to predict consensus positions that differ from the target sequence (Figure 5).

Our third finding is that these trends do not directly correlate with data volume. Visual inspection indicates that training performance declined from 2015 to 2019 (Figure 2E), then significantly improved in 2022 with just one year’s difference. Upon inspection, we found that these datasets do not represent a continuous growth of valid observations; they change both by adding new data and by culling some older data. UniRef data is routinely removed when found to be invalid, and in 2021, there was an unusually large cull – twice the usual amount that significantly improved the training quality of the data (Figure 2F). Taken together, these findings suggest that pLMs should be retrained on a regular basis.

### The Computational Costs of pLMs Can Substantially Be Reduced

The findings in the previous section have significant implications for the literature, as models trained in recent years are often compared directly to state-of-the-art models that were trained without the benefit of improved data quality. In this work, we addressed this phenomenon by training our own ESM2 controls on the newest datasets. However, these measures were only possible because of access to open-source code for curating datasets and for efficient pretraining of pLMs. Such open-source access is thus necessary in order to ensure that the scientific community can fairly evaluate the latest models on the latest datasets.

As detailed in the Methods section, we have adopted several recent advances in training large language models to increase computational efficiency, such as the adoption of FlashAttention and our handling of long sequences. Those improvements, combined with the previously described benefits of curating the training and test data to enable smaller model sizes, dramatically reduced the costs of training effective models compared to existing codebases and data pipelines. Following a popular technique for estimating the number of FLOPs to train LLMs [Hoffmann et al., 2022], AMPLIFY 120M required 1.6×10^21^ FLOPs to train for 1M steps while being comparable to ESM2 3B, which requires 11× more theoretical compute at 1.8×10^22^ FLOPs. Similarly, AMPLIFY 350M requires 4.6×10^21^ FLOPs while outperforming ESM2 15B, which requires 17× more compute, at 8.1×10^22^ FLOPs. However, theoretical FLOPs do not tell the whole story. While ESM2 150M only requires 1.25× more FLOPs per training step compared to AMPLIFY 120M, our implementation of AMPLIFY 120M trains 2× faster than the HuggingFace implementation, even with optimizations such as compilation and mixed-precision, further democratizing the training of state-of-the-art pLMs.

Beyond training costs, our models are an order of magnitude more efficient during inference than any ESM2 models of comparable performance (Figure 1C). This is both because we attain equivalent performance with significantly smaller models, and because the incorporation of FlashAttention lightly improves the scaling behavior observed for sequence length. The ESM2 15B model approaches the performance of our 350M parameter model, but at an inference cost upwards of 24× to 29× higher, depending on sequence length (Figure 1C). Such differences during inference are critical when trained models are deployed for large-scale in silico screens of protein sequences, such as are performed during the design of novel proteins with prescribed properties.

### AMPLIFY Exhibits Zero-Shot Emergent Behaviors

One of the most intriguing aspects of pLMs is their ability to exhibit emergent behaviors that can be used to study or predict protein properties, despite being trained in an unsupervised manner. In AMPLIFY, we have observed several such behaviors, including the ability to identify sequences that may not be proteins and to distinguish these from natively disordered proteins, which are known to have different statistical properties from ordered proteins. Figures 2A-C show UMAP projections of AMPLIFY’s residue embeddings specifically for the arginine position of RG motifs in human proteins, chosen here to represent a common sequence motif with clearly defined function [Thandapani et al., 2013]. The tightly clustered regions in the upper right and lower left of these plots can be used to highlight some of AMPLIFY’s emergent properties.

Figure 2A is color-coded by AlphaFold2’s confidence in predicting the structure of the corresponding proteins. Low-confidence predictions can indicate gaps in AlphaFold2’s training data (i.e., the PDB, which is much smaller than sequence databases) or natively disordered proteins. These low-confidence predictions are spread throughout the space but are especially concentrated in tightly clustered regions. Further analysis shows that most known natively disordered proteins fall outside these regions, suggesting that these clusters may be enriched for proteins that span gaps in AlphaFold2’s training data.

Figure 2B is color-coded by three gene ontology (GO) term labels [Ashburner et al., 2000] to reveal aspects of the space. The lower left region, where most of the sequences are colored green, is enriched for the collagen trimer GO term. The upper right region, where many of the sequences are colored magenta, is enriched for proteins without any GO annotation for protein localization at all, lacking even an annotation for unknown localization. The lack of any annotation data despite an often 20+ year inclusion in the reference human proteome suggests these sequences may not be proteins at all. The blue-colored sequences have the RNA Binding GO annotation, demonstrating that AMPLIFY does not always cluster sequences by their GO annotations or lack thereof.

Figure 2C is color-coded in three ways. Blue circles and magenta diamonds both represent sequences with the lowest amount of evidence (PE=5, uncertain hypothetical proteins) according to UniProtKB. This category includes high-risk sequences that may not actually be proteins. However, blue circles have GO annotations for subcellular localization, whereas magenta diamonds do not. Intuitively, a protein with a localization annotation is more likely to be real, despite having a PE=5 annotation than one that does not. Consistent with this intuition, the blue circles are more evenly spread throughout the UMAP space than the magenta diamonds are. Closer examination reveals that some of these latter sequences have recently been annotated as originating from long non-coding RNAs and can be classified as confirmed non-proteins [Mattick et al., 2023].

These observations suggest that the upper right region of the learned space corresponds to where the model places sequences it does not ‘recognize’ as real proteins. We tested this hypothesis in two ways. First, we embedded a gold standard non-protein sequence – the first chapter of Mary Shelley’s Frankenstein [Shelley, 2018]. As shown by the yellow stars in Figure 2C, this sequence localizes to the same upper right portion of the space as the magenta diamonds.

The second test involved comparing all residue embeddings for human proteins with PE=1 (evidence at protein level) against those with PE=5, considering all residues across each sequence. Figure 2D shows that, similar to Figure 2C, PE=5 sequences are, on average, very similar to the embedding of the first chapter of Frankenstein. Conversely, PE=1 sequences are almost entirely dissimilar to Frankenstein. Within each sequence we also measure similarity between individual residue embeddings and the sequence mean embedding and find that PE=5 sequences become similar by converging.

Both similarity to Frankenstein and embedding convergence are highly predictive for discriminating between PE=1 and PE=5 in the human proteome (Figure 2D). We then revisited the experiment on UniRef100 data quality over time, scoring all sequences included in UniRef100 since 2011 based on their similarity to our non-protein test. This analysis revealed that the sequence cull in 2021, which improved model training performance in 2022, was not only larger than in other years but was also specifically enriched for sequences that might not be proteins at all (Figure 2F).

Suspecting that the proportion of these questionable proteins in the training data consistently affects model quality, we revisited why training on UniRef100 improves model quality compared to UniRef50. Analyzing the embeddings for all proteins in both sets, we found that UniRef100’s sequence distribution closely mirrors the natural distribution in the human proteome. In contrast, clustering has shifted UniRef50 significantly towards a space that may not represent real proteins (Figure 2G).

ESM demonstrated that attention patterns within pLMs correlate with structural contacts. Interestingly, while AMPLIFY outperforms even the best ESM2 on sequence recovery, ESM2 outperforms AMPLIFY at the task of unsupervised contact prediction (Table 8). A key distinction of AMPLIFY’s architecture compared to ESM2 is its smaller number of attention heads. Since ESM’s methodology relies on these heads for contact prediction and has more of them, AMPLIFY may be at a disadvantage. As a result, structural information might not emerge as clearly in the attention maps in a linear fashion. To explore this hypothesis, we replace the logistic regression with a random forest. We find that the performance gap on unsupervised contact prediction decreases substantially by using the non-linear classifier (Table 9).

**Table 8:** ESM style contact prediction. The standard deviation is computed across the proteins in each dataset.

| Architecture | #Parameters | #Heads | CASP14 |  | CAMEO |  | CASP15 |
| --- | --- | --- | --- | --- | --- | --- | --- |
|  |  |  | Ours | Reported | Ours (Jan-Mar 24) | Reported (Apr-Jun 22) | Ours |
| ESM2 | 8M | 120 | 8.9 ± 8.8 | 9.8 | 12.4 ± 12.7 | 15.7 | 11.4 ± 12.5 |
| | 35M | 240 | 14.5 ± 12.5 | 16.4 | 19.3 ± 15.6 | 28.4 | 22.5 ± 13.6 |
| | 150M | 600 | 25.5 ± 18.7 | 26.8 | 26.8 ± 17.7 | 40.1 | 30.5 ± 16.1 |
| | 650M | 666 | 31.6 ± 24.3 | 32.5 | 34.4 ± 20.0 | 47.6 | 35.9 ± 17.6 |
| | 3B | 1,440 | 35.5 ± 24.5 | 34.0 | 34.8 ± 19.1 | 49.9 | 36.8 ± 17.6 |
| AMPLIFY | 120M | 240 | 12.4 ± 11.3 | | 17.8 ± 14.1 | | 16.9 ± 13.2 |
| | 350M | 480 | 16.6 ± 13.6 | | 20.9 ± 15.7 | | 20.0 ± 14.6 |

**Table 9:** Random forest contact prediction. The standard deviation is computed across the proteins in each dataset.

| Architecture | #Parameters | #Heads | CASP14 | CAMEO | CASP15 |
| --- | --- | --- | --- | --- | --- |
|  |  |  | Ours | Ours (Jan-Mar 24) | Ours |
| ESM2 | 8M | 120 | 36.4 ± 39.8 | 11.5 ± 11.2 | 48.5 ± 37.2 |
| | 35M | 240 | 39.5 ± 39.1 | 19.6 ± 15.5 | 52.9 ± 33.9 |
| | 150M | 600 | 44.3 ± 38.3 | 26.7 ± 17.8 | 55.8 ± 31.7 |
| | 650M | 666 | 45.9 ± 38.2 | 34.0 ± 19.5 | 58.3 ± 30.1 |
| | 3B | 1,440 | 48.1 ± 36.9 | 34.6 ± 18.9 | 58.2 ± 29.7 |
| | 15B | 1,920 | 48.7 ± 36.8 | 35.3 ± 19.1 | 58.8 ± 29.8 |
| AMPLIFY | 120M | 240 | 39.4 ± 39.1 | 17.6 ± 13.6 | 51.0 ± 34.0 |
| | 350M | 480 | 40.9 ± 39.3 | 21.0 ± 16.1 | 52.4 ± 32.7 |

### Comparison to Structure-Based Models

Like other pLMs, AMPLIFY also exhibit the ability to learn structural features in an unsupervised fashion from sequence data alone. In particular, embedding similarity serves as a zero-shot predictor for various structural properties, and attention patterns correlate with structural contacts (Table 9). While models such as AlphaFold2 which are specifically trained from structure can predict these properties better, there is a clear advantage to training on natural sequence space without available structural data. As previously mentioned, AlphaFold2’s confidence has been used to predict protein disorder [Ruff and Pappu, 2021], but we show that AlphaFold2 cannot differentiate between disordered proteins and non-protein sequences, whereas AMPLIFY can. Figure 2H demonstrates that AMPLIFY embedding similarity can separate human disordered proteins (>25% intrinsically disordered in the DisProt database) [Aspromonte et al., 2024] from hypothetical proteins (PE=5 and no annotation for localization) at a ROC-AUC of 0.94, compared to a score of 0.44 when using AlphaFold2’s pLDDT confidence metric. This demonstrates that training solely on available structures is insufficient for a comprehensive understanding of protein behavior.

A common pattern in computational protein design is to predict the structure of a designed sequence and then to predict new sequences for that structure. For this purpose the pairing of AlphaFold2 and ProteinMPNN models has been shown to be highly effective, especially for the task of salvaging structure-based designs that fail to fold or are insoluble [Dauparas et al., 2022]. We benchmark this category of methods against our human proteome sequence recovery tasks and show that for natural human proteins sequence-to-structure-to-sequence methods perform poorly at both sequence recovery and the ability to predict most common non-target residues (Figure 9).

## Discussion

In recent years, there has been a growing trend in machine learning to scale neural networks, particularly language models for text or proteins, taking advantage of the performance benefits that come with scale [Kaplan et al., 2020, Hoffmann et al., 2022]. This has led to a situation where the training costs of best-in-class models are often out of reach for most of the scientific community. As a result, the largest and most effective models are trained by private institutional organizations and then released with restrictions in order to recoup their investment. However, our study shows that this is not necessary. Instead, the expertise of protein scientists can be leveraged to curate the training task to ask the right questions, which is competitive with scale in terms of performance while being significantly more efficient.

The curation of protein data for machine learning is a tricky task. We are using machine learning to improve our understanding of proteins, yet a better understanding of proteins is precisely what is needed to improve data curation. Human understanding of proteins is very limited compared to that of natural language, as our findings show that even an expert can struggle to identify whether a given protein sequence is even real.

The role of expertise in building efficient models makes it essential to define expertise. As authors, we must emphasize that the expertise we found to be competitive with scale was not our own, but rather the collective expertise of the broad community of protein scientists producing, studying, and annotating the data in the pursuit of our shared goal. Our work stands on this expertise.

A key finding from our analysis is the potential application of machine learning to identify protein sequences that should be reevaluated by experts, as the pLM training process cannot explain them. By releasing our methods and data as open-source, we aim to support a diverse set of researchers working on fundamental problems in protein science and promote iterative rounds of data curation and model building. We depend on their advances to develop new therapies for patients.

## Methods

### Construction of the Test Data Sequence Sets

In total, three protein sequence test sets were constructed: reference proteomes sequences from UniProt, paired-antibodies from the Observed Antibody Space (OAS) database [Kovaltsuk et al., 2018], and single domains from the Structural Classification of Proteins version 2 (SCOP 2) database [Chandonia et al., 2022] to allow task-specific validation of the models. UniProt represents the natural distribution of proteins and is the main test set. OAS antibody sequences were included to allow assessing the performance on complementarity-determining regions (CDR) loops of antibody sequences and SCOP protein sequences were included to allow sequence-to-structure specific tasks.

Reference proteomes derived from high quality genome assemblies (BUSCO completeness score >80%) [Simão et al., 2015, Seppey et al., 2019] were selected as source data to construct the reference proteome sequence test dataset. Reference proteomes of organisms covering all three domains of the phylogenetic tree of life were included (Bacteria, Archaea and Eukarya). Next, any protein sequence lacking experimental evidence at the protein or transcript level (protein existence (PE) level 1 & 2) or that were comprised of 5 or less amino acids were removed from the dataset. Paired heavy and light chain antibody sequences were retrieved from the OAS database and the SCOP family representative domain sequences were retrieved from SCOP. From all sets, protein sequences that contained non-canonical, rare, or ambiguous amino acids (B, J, O, U, X, or Z) were removed. Next, sequences from the OAS set and the SCOP set were clustered using MMseqs2 [Steinegger and Söding, 2017] at 90% and 30% sequence identity, respectively, and only the representative sequence for each cluster was retained. Importantly, the UniProt test set was not clustered to preserve the natural distribution of proteins. From the SCOP and the OAS set, 10K protein sequences were randomly selected from each and used in the SCOP and OAS test sets, respectively. Highly similar protein sequences in the reference proteome set (>90% sequence identity) with the OAS and SCOP test sets were removed, and, finally 10K randomly selected reference proteome sequences were selected as the reference proteome dataset.

### Construction of Training Data Sequence Sets

UniRef, OAS, SCOP, and UniProt databases were downloaded in December 2023 to use as primary basis sets for test and training. All data was processed by removing sequences with rare or ambiguous amino acids (B, J, O, U, X, and Z) and by using MMseqs2 to exclude sequences with >90% sequence identity to sequences in our test sets. Antibodies from OAS were excluded at a sequence identity of >99% to account for the high similarity across germline frameworks, as we are primarily interested in minor variations within CDR loop sequences. To create our combined OAS, SCOP, and UniRef datasets we subselected from OAS only the paired sequence data, augmenting the dataset by incorporating heavy (Hc) and light (Lc) chain sequences separated by a chainbreak token ‘|’, in both Hc|Lc and Lc|Hc orderings.

### Model

To increase training and inference efficiency, we incorporated recent improvements in LLMs for NLP [Touvron et al., 2023]. Several key modifications distinguish AMPLIFY from ESM2. Architecturally, we replaced the GeLU activation function and LayerNorm with a fused implementation of SwiGLU [Shazeer, 2020] and RMSNorm [Zhang and Sennrich, 2019], respectively, and adjusted the depth-to-width ratio based on the theoretical analysis of Levine et al. [2020]. Additionally, the model employs a fast and memory-efficient exact attention mechanism to significantly reduce the computational cost of training [Dao et al., 2022]. We also reduced the number of attention heads so that each head has a dimension of 64, enhancing their representation power.

At the optimization level, we replaced the Adam optimizer with the improved AdamW [Loshchilov, 2017], using tuned beta hyperparameters and a higher learning rate. The linear scheduler was swapped for a cosine annealing scheduler, and the number of gradient steps was increased from 500K to 1 million. The model was trained with lower precision (bf16), utilizing DeepSpeed [Aminabadi et al., 2022], and model sharding [Rajbhandari et al., 2020]. We also enabled gradient clipping set to 1 for all models.

We streamlined the vocabulary by removing unused tokens like O,.,-, and <null_1>, reducing its size from 33 to 27. Additionally, we added a new token ‘|’ to serve as a chain separator for multi-chain proteins, such as antibody heavy and light chains. In the training set, 80% of the sequences were 512 residues or shorter. To improve performance, sequences longer than 512 amino acids were randomly truncated during training to minimize padding. This truncation alters the natural distribution of sequence lengths, so after training on truncated sequences, we expanded the context to 2048 residues and trained AMPLIFY 120M and 350M for an additional 25K and 50K steps, respectively (Table 3 and Figure 6).

We also considered and evaluated several other modifications on a smaller scale, including label smoothing, class weighting, span masking, excluding special tokens from replacement, and learning rate warm restarts. However, these did not yield promising results, so they were not tested on a larger scale and are not reported here.

### Retraining ESM2

To test the training performance of ESM2 models, we established a control where HuggingFace is used to retrain ESM2 architectures using updated versions of UniRef and our validation criteria.

### Perplexity and Pseudo-Perplexity

We used the same definitions for perplexity and pseudo-perplexity as ESM. Perplexity is computed on the 15% masked tokens. Pseudo-perplexity is computed by masking each residue one at a time. While more expensive, the pseudo-perplexity does not suffer from the stochasticity where one may mask residues that are ‘easy’ to recover and therefore overestimate the model’s performance.

## Acknowledgements

The authors would like to thank Amgen’s Alan Russell, Marti Head, Peter Grandsard, Carolyn Ch’ing, Matt Foster, and Karen Chou, Mila’s Robert Farias, Lola Le Breton, Joseph Viviano, and Mathieu Bourgey, and NVIDIA’s Alex Sabatier, David Klien, Sam Kjellesvig, and Greg Zynda.

Sarath Chandar is supported by the Canada CIFAR AI Chairs program, the Canada Research Chair in Lifelong Machine Learning, and the NSERC Discovery Grant. The project was also supported by the Amgen-Mila collaboration grant. The authors acknowledge the computational resources provided by Amgen and Mila.

## Data Availability Statement

Our dataset includes proteins from UniProt [Consortium, 2022], the Observed Antibody Space (OAS) [Kovaltsuk et al., 2018], and individual domains from the Structural Classification of Proteins (SCOP) [Chandonia et al., 2022] databases. We released the entire dataset on Zenodo^3^ and Hugging Face^4^ under an MIT license. Zenodo is an open-source, open-access, and open-data platform launched by CERN in 2013. Hugging Face is a popular platform for building machine learning applications.

## Code Availability Statement

We released the entire codebase on GitHub^5^ and the models on Hugging Face^6^ under an MIT license. GitHub, a Microsoft subsidiary, is a platform for code creation, management, and collaboration. Hugging Face is a widely popular platform for building machine learning applications.

## Contributions

C.J.L. and S.C. led the research. R.V. and A.v.d.S. developed and collected the data. Q.F. developed the neural network architecture and training. Q.F. and R.V. designed the experiments. Q.F., R.V., and B.S. conducted the experiments. B.S. contributed technical advice and ideas. B.S. and Q.F. prepared the code for open source release. All authors wrote the manuscript.

## AMPLIFY

### Architecture

AMPLIFY is a Transformer encoder based on BERT [Devlin et al., 2019] that incorporates widely adopted modifications from modern large language models, such as Llama2 [Touvron et al., 2023]. Layer normalization is applied before the self-attention and feed-forward blocks (PreLN), which has been shown to stabilize training [Xiong et al., 2020]. Additionally, layer normalization is replaced with root mean square layer normalization (RMSNorm), offering comparable performance at a lower computational cost [Zhang and Sennrich, 2019]. Positional encoding is substituted with Rotary Position Embedding (RoPE) [Su et al., 2024], which encodes absolute positions using a rotation matrix directly added to the queries and keys, eliminating the need for the Transformer to preserve positional information across layers. The GeLU activation function is replaced with SwiGLU [Shazeer, 2020], a gated linear unit (GLU) that computes the element-wise product of two input projections, one of which passes through the Swish activation function. Given the relatively small size of the network, dropout is omitted to preserve the model’s expressive power. Moreover, the number of layers and their width are tuned to achieve the optimal depth-to-width ratio following the theoretical analysis of Levine et al. [2020], maximizing the network’s representation power. The number of attention heads is reduced to ensure each head has a dimension of 64, providing sufficient representation power. Finally, all biases are removed except those in the output classification head. The configuration details for AMPLIFY 120M and 350M are given in Table 1.

### Training

AMPLIFY is trained with the masked language modeling (MLM) task, where 15% of the residues are randomly selected, of which 80% are masked, 10% are randomly replaced, and 10% are left unchanged [Devlin et al., 2019]. The model is trained using cross-entropy loss and optimized with AdamW. The learning rate is first linearly increased from 0 to the peak learning rate during the warmup phase before being decayed using a cosine learning rate scheduler over 90% of the gradient steps. AMPLIFY is first trained for 1 million steps on sequences truncated to 512 residues for efficiency, then extended for a small number of steps to 2,048 residues. Special beginning-of-sequence (BOS) and end-of-sequence (EOS) tokens are added around the proteins prior to truncation, enabling the model to differentiate between full-length and truncated proteins. The training details are given in Table 3.

### Efficiency

AMPLIFY 350M outperforms the best sequence-only pLM, ESM2 15B, with 43× fewer parameters while requiring 17× fewer FLOPs to train and achieving 24 to 29× higher throughput at inference time, depending on sequence length. Several factors explain such a drastic speed-up.

AMPLIFY exhibits greater representation power than an equivalent-sized ESM2, thanks to the architectural modifications previously described. Additionally, our implementation leverages fused operators from the xFormers library, and the hidden and head dimensions were chosen to be multiples of 64 to optimize the usage on modern GPUs. Biases are removed from both the feedforward network and self-attention layers to simplify the computation. Lastly, the memory demands of self-attention’s quadratic complexity are mitigated with FlashAttention, a fast and memory-efficient exact attention. Although the computational complexity remains the same, the n-square matrix is not stored in memory, significantly improving speed scalability with sequence length.

AMPLIFY is trained using mixed-precision (bf16) [Micikevicius et al., 2017], which allows modern GPUs to achieve 2× FLOPs compared to full precision (fp32). The model is distributed across machines and parallelized across devices with DeepSpeed [Aminabadi et al., 2022]. The parameters, activations, and gradients are not duplicated across GPUs thanks to the Zero Redundancy Optimizer (ZeRO) [Rajbhandari et al., 2020], reducing memory usage compared to the naive distribution approach and enabling larger batch sizes per device, thereby fully leveraging the highly parallel nature of GPUs.

HuggingFace does not benefit from all of these implementation enhancements, resulting in significantly slower performance. Even with mixed-precision (bf16) and JIT-compiling PyTorch code into optimized kernels, applying similar though less extensive optimizations as fused techniques, HuggingFace models lag behind in terms of efficiency. For example, training AMPLIFY 120M is twice as fast per step as ESM2 150M from HuggingFace. While both models took approximately 200 actual A100 GPU days, the theoretical cost of ESM2 150M is 75 A100 days, while AMPLIFY is 119 A100 days. This indicates that Hugging Face’s overhead is more than 1.5× greater than ours. At inference, padding is omitted when computing the pseudo-perplexity as each residue is masked individually, creating batches of same-size sequences. The xFormers library is able to leverage a more efficient implementation of FlashAttention when no padding or dropout is used, further increasing the throughput and scaling more effectively with sequence length. This optimization is impossible during training because proteins with various lengths are packed together and, therefore, require padding. The most significant speed-up is observed when comparing AMPLIFY 350M to its closest competitor, ESM2 15B. On a single A100, AMPLIFY can evaluate 584.7 ± 0.5 sequences of length 256, whereas ESM2 15B can only manage 21.3 ± 0.0 sequences. The relative performance gap widens as sequence length increases thanks to FlashAttention, reaching its peak with sequences of 1,024 residues, where AMPLIFY achieves a throughput of 135.7 ± 0.1 compared to ESM2’s 4.7 ± 0.0. Performing inference on one sequence at a time narrows the gap between the models, as AMPLIFY cannot leverage larger batch sizes. At a sequence of 1,024, AMPLIFY 350M has a throughput of 34.6 ± 0.2, while ESM2 15B has a throughput of 4.6 ± 0.0. The throughput of both models is provided in Table 4.

The theoretical number of operations required to train a model can be calculated independently of the implementation, excluding overhead factors like GPU communication, checkpointing, and evaluation. AMPLIFY 350M has a theoretical cost of 4.92×10^21^ FLOPs, whereas ESM2 15B requires 8.09×10^22^ FLOPs. Assuming all operations are performed using TF32, an A100 GPU has a theoretical throughput of 156 TFLOPs^7^; thus, AMPLIFY 350M theoretically requires 365 A100 GPU days, while ESM2 15B demands 6,000 A100 GPU days. However, implementation plays a crucial role in practice. AMPLIFY 350M required 1,014 actual A100 GPU days to train. While we do not have access to the exact implementation Meta used for ESM2 15B, the authors estimated that it required 30,720 V100 GPU days^8^. AWS offers on-demand instances with 8 x A100 for $40.96 per hour^9^ and 8 x V100 for $31.218 per hour^10^. Based on these rates, training AMPLIFY 350M would cost $124,600, while training ESM2 15B would cost $2,877,050. However, this comparison is unfair to ESM2, as the V100 is an older model than the A100. According to Nvidia’s documentation, for FP16/FP32 mixed-precision, A100 delivers 2.5× the performance of V100^11^. Therefore, we estimate that ESM2 15B would have required at least 12,000 A100 GPU days to train, bringing its cost on A100 to approximately $1,474,560.^^

### Effect of Clustering on Training Data

The data observed during training defines the fitness landscape that models are intended to learn. For foundation models, which excel at capturing complex distributions, it is essential that this data accurately represent the unknown real-world fitness landscape. Therefore, any systematic biases in how data is observed and deposited must be addressed.

One of the most common biases in biological sequence data is redundancy. Some sequences get measured and deposited multiple times, increasing their prevalence without representing novel observations. Typically, redundancy is addressed by clustering sequences and filtering them for novelty. However, this collapses novel observations into a single data point, for instance when two species share a protein sequence. Despite this flaw, filtering for novelty is still widely used as an efficient and self-contained redundancy correction method.

The UniRef project tackles redundancy by filtering in this way, resulting in UniRef100. Additionally, the UniRef project also establishes datasets based on clustering thresholds of 90% and 50% sequence identity, known as UniRef90 and UniRef50, respectively. It is important to note that these datasets were not originally designed to counteract redundancy bias but rather to identify which novel observations in UniRef100 are most likely to share biological functions with one another.

In recent years many research programs have instead treated these lower identity cutoffs as stricter handling of redundancy bias, and UniRef50 is often presented as the least biased dataset. As a result many methods end up training exclusively on UniRef50. Even ESM2, which used a larger population of sequences for training, relied on UniRef50 to establish sampling distributions and test sets.

There is a fundamental problem with clustering. Outside of exact matches, identity statistics are predictions made by sequence alignment methods, and these methods introduce systematic biases depending on the difficulty of aligning different categories of protein sequences. Proteins with functions that require significant positional conservation, such as individual folded domains, are easy to cluster. In contrast, proteins that do not have positional conservation, including disordered proteins, tandem repeat proteins, and longer proteins where domains can swap places without affecting function, are more challenging to cluster. This is also true, quite critically, for sequences that do not require positional conservation because they are, in fact, not protein sequences at all. This includes pseudogenes, incorrect gene predictions, and non-coding RNA that has been translated into protein sequences and deposited effectively in error.

By using unclustered test sets, we demonstrate that UniRef50 impairs model training. The nature of the bias itself is worth describing, and we give some examples of this bias in Table 5.

For the subset of UniRef proteins that come from the UniProt-SPROT and TrEMBL datasets, we can do simple enrichment analysis for keywords and metadata stored in the protein name field of the source FASTA files, demonstrating that the clustering step introduces clear bias to the sequence distribution that up-weights proteins expected to have low positional conservation and down weights proteins with high conservation.

For protein evidence levels, the bulk of sequences in both UniRef sets come from predictions that are either inferred from homology (PE=3) or are a more general prediction (PE=4). In UniRef100, these exist in a ratio of 1.8, but removing sequences by clustering is more likely to affect proteins with clear homologs. In UniRef50, this ratio increases to 6.1, skewing the distribution dramatically towards lower confidence sequences.

For keywords, the protein name terms that are enriched in UniRef50 include many categories where we expect sequence alignment-based identity matching to have a harder time tracking functional relationships. This includes repeat proteins and proteins that are listed as containing a specific known domain, which is a phrase that usually means there’s a high likelihood of multiple domains or disordered elements. We also see enrichment in terms that have a higher risk of not being true protein sequences, with proteins that have ‘hypothetical protein’, ‘uncharacterized’, and ‘intergenic region’ in their assigned names.

On the other side, UniRef50 specifically depletes the sampling of proteins with known functions and a general expectation of strong positional conservation. This includes named kinases, polymerases, receptors, enzymes, transporters, ATP-binding proteins, and chaperones. Chaperones show a remarkable five-fold depletion, with 153,827 sequences pared down to only 7,712. Protein chaperones are some of the most conserved proteins in evolution, and biologists use them to establish relationships between organisms that diverged over a billion years ago. Reducing their weight during training by labeling that conservation as redundancy is a clear mistake.

### Orthogonal Testing

#### Human Proteome Test Set

When using pLMs for protein design, the objective is not to replicate the input sequence but to identify functional sequences that differ from it. This task is fundamentally distinct from the mask replacement task we train on and requires the development of separate evaluation methods.

To evaluate sequence behavior beyond simple mask replacement, we selected the human proteome (December 2023) as a representative sequence set. We then constructed sequence profiles using the MSA method outlined by Jumper et al. [2021], focusing on a final set of 6,823 proteins. This selection was based on the following criteria: sequence length between 1 and 1024, no ambiguous amino acids, MSA with at least 50 homologs, and sequences used either in training or testing (excluding proteins that were omitted from the training set only due to similarity to the test set). This approach captures 2,599,705 residue positions, which we categorize into 8 discrete, non-overlapping groups based on observed frequencies.

Highly conserved residues are preserved through evolution due to their functional importance. The aim of pLM training is not merely to reconstruct existing sequences but to understand and generalize the underlying relationships that contribute to function prediction. From a sequence recovery perspective, the higher the conservation, the more crucial it is for pLMs to correctly identify conserved residues. This holds true whether the conserved identity matches the starting sequence or differs from it.

^11^Out of memory.

To capture this, we’ve divided residues into 8 categories based on both the conservation of the human residue (the target) and the conservation of the most common non-target residue. The specific conservation frequencies were chosen to ensure that each category has a sufficient sample size, even when further subdividing by training set, test set, and sequence length from 1 to 510 or 511 to 1024. The categories and the number of residues in each are summarized in Table 6.

When working with this set, we identify Category B as the most reliable test for positions where the target identity is highly conserved, making mask replacement the ideal task, followed by Categories A and C. Although Category A has a higher frequency cutoff than B, sequencing errors mean we don’t expect 100% conservation, and achieving it often indicates that very few homologs have been identified for that position.

Category I serves as the best orthogonal test, capturing positions where the target human sequence is not conserved but an alternative residue identity is highly conserved. Categories H, F, and E are similar in this respect but with a smaller difference in conservation between the target and non-target identities.

Categories D and G represent positions where no single residue is conserved across the majority of sequences. It is unclear what the ideal behavior of a pLM should be at these positions, as they either lack conservation or have a conservation profile better described by sets of synonymous residues rather than individual identities.

To evaluate model performance on these sequence categories, we first use all models in a mask replacement mode to predict sequence logits. We then test the highest-scoring amino acid token against the real sequence, measuring the frequency of recovering both the original masked residue (the’target residue’) and, in cases where a non-target amino acid is more common in homologs, the frequency of recovering this alternative residue (the’most common non-target residue’).

For residues in highly conserved positions where mask replacement is clearly the appropriate task (Category B), model accuracy reaches upwards of 99% (for AMPLIFY 350M, as shown in Figures 3 and 4). However, when conservation decreases (Category C), this recovery rate also declines, with a maximum accuracy across all models in the 92-94% range. Given the high accuracy in Category B, we hypothesize that this decline reflects not a decrease in model accuracy but a decrease in metric accuracy. We believe these mismatches often represent predictions of valid synonymous residues.

**Figure 3:**
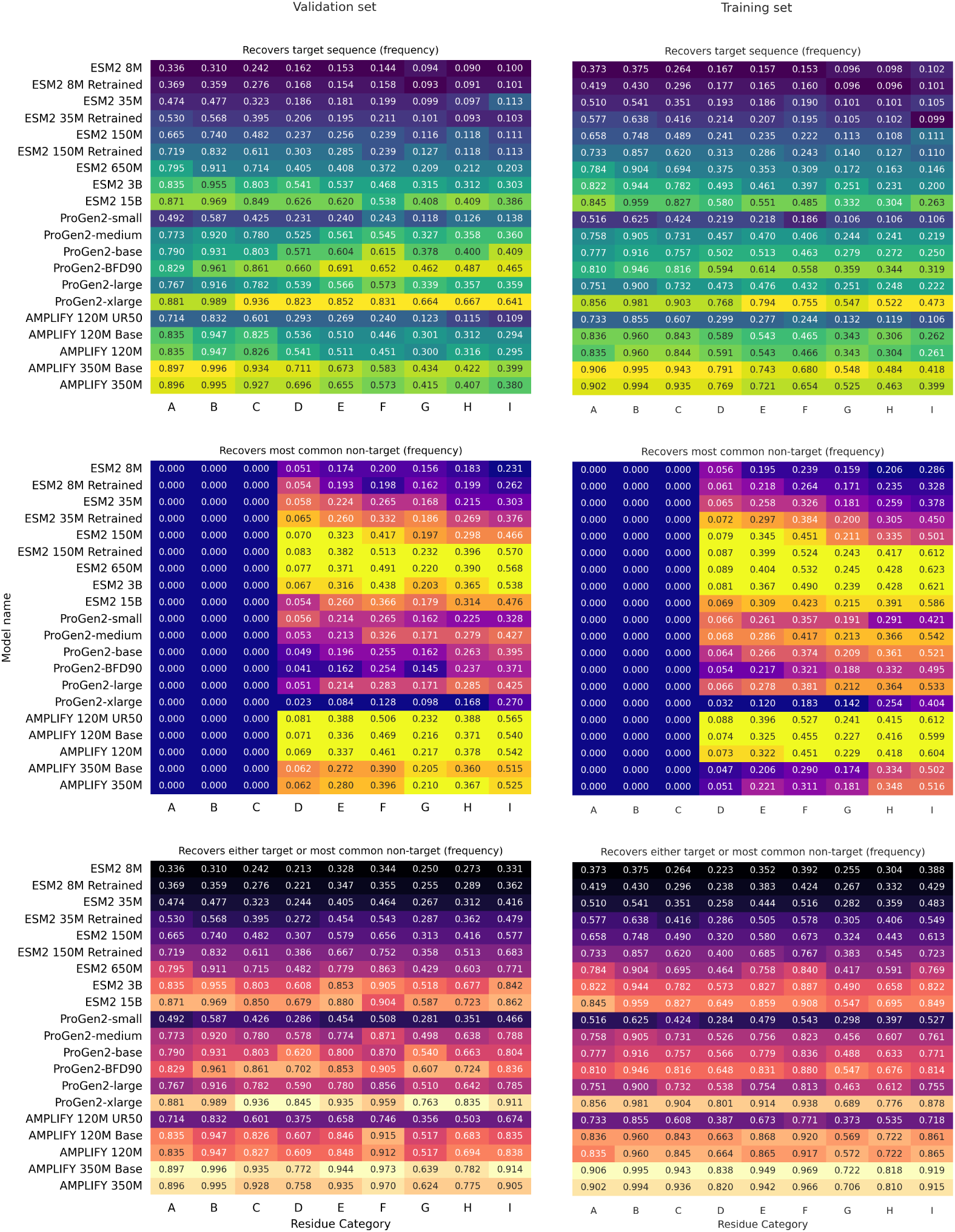
Sequence recovery for lengths from 1 to 510 as a function of the model and category. Residue recovery rates for a selection of models vs. human test set residue categories as defined in Table 6, and covering sequence lengths within the context limit of primary training (1-510 residues). The top row of panels shows rates for target sequence recovery, middle panel shows rates for recovering the most common non-target sequence, and bottom panels show the combined rate of recovery for either target sequence or most common non-target. Rates for test set proteins (not seen during AMPLIFY training) are shown on the left, and rates for training set proteins are shown on the right.

**Figure 4:**
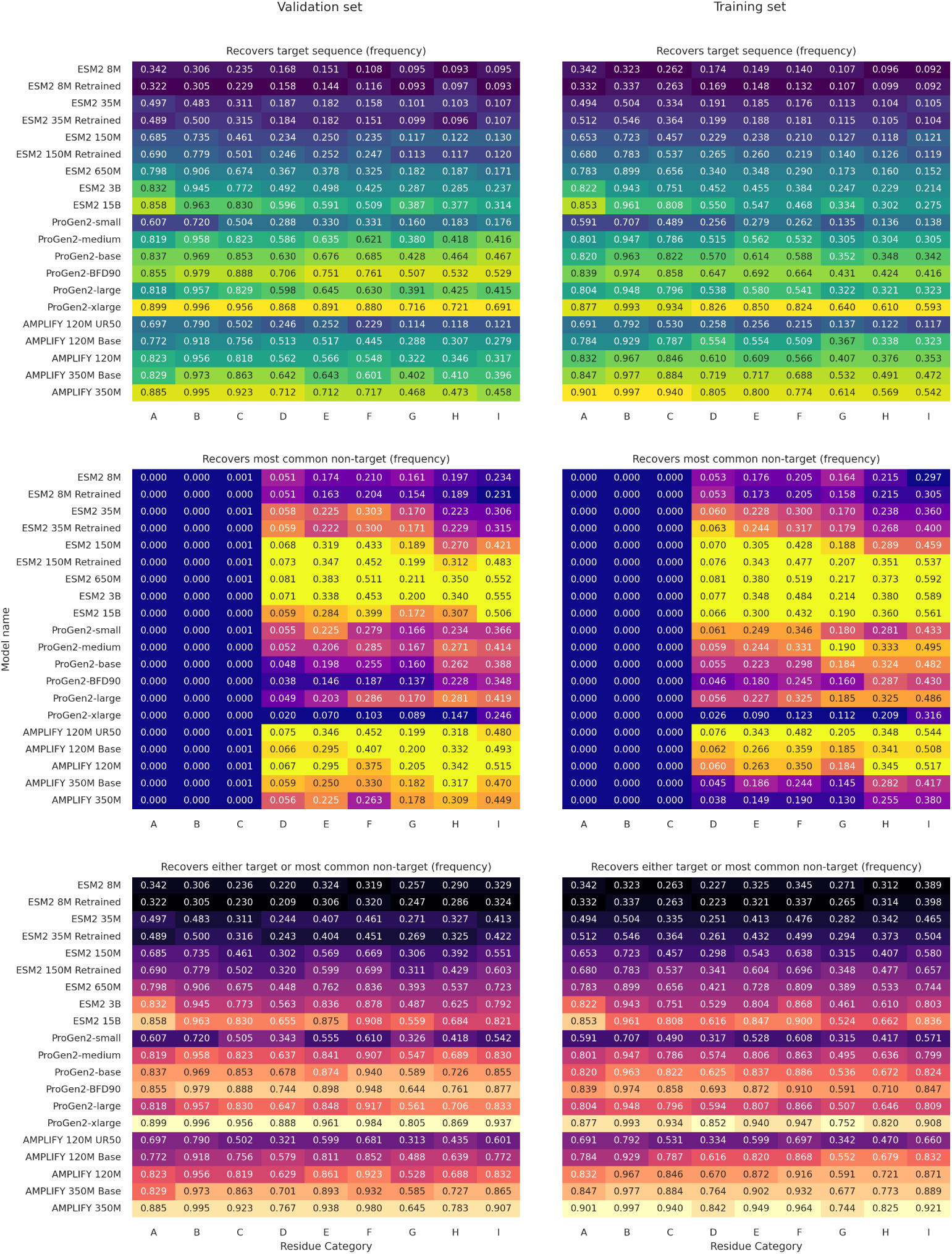
Sequence recovery for lengths from 511 to 1024 as a function of the model and category. Residue recovery rates for a selection of models vs. human test set residue categories as defined in Table 6, and covering longer sequence lengths only seen in their entirety during context extension (511-1024 residues). The top row of panels shows rates for target sequence recovery, middle panel shows rates for recovering the most common non-target sequence, and bottom panels show the combined rate of recovery for either target sequence or most common non-target. Rates for test set proteins (not seen during AMPLIFY training) are shown on the left, and rates for training set proteins are shown on the right.

**Figure 5:**
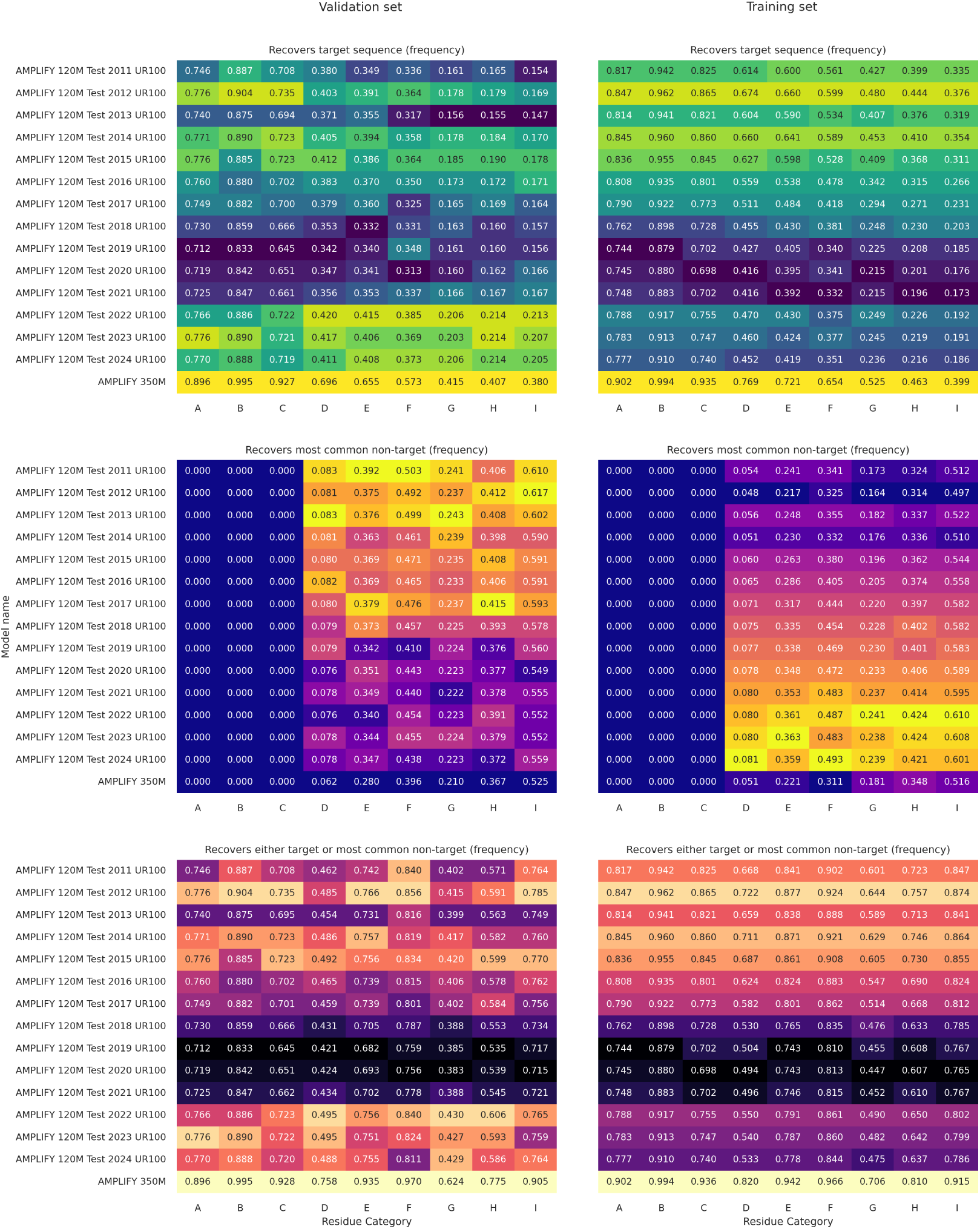
Sequence recovery for lengths from 1 to 511 as a function of the UniRef yearly release and category. Residue recovery rates for test 120M parameter models trained on yearly UniRef releases spanning 2011 to 2024, with AMPLIFY350M presented as a control. Rates are shown for test set residue categories as defined in Table 6, covering sequence lengths within the context limit of primary training (1-510 residues). The top row of panels shows rates for target sequence recovery, middle panel shows rates for recovering the most common non-target sequence, and bottom panels show the combined rate of recovery for either target sequence or most common non-target. Rates for test set proteins (not seen during AMPLIFY training) are shown on the left, and rates for training set proteins are shown on the right.

**Figure 6:**
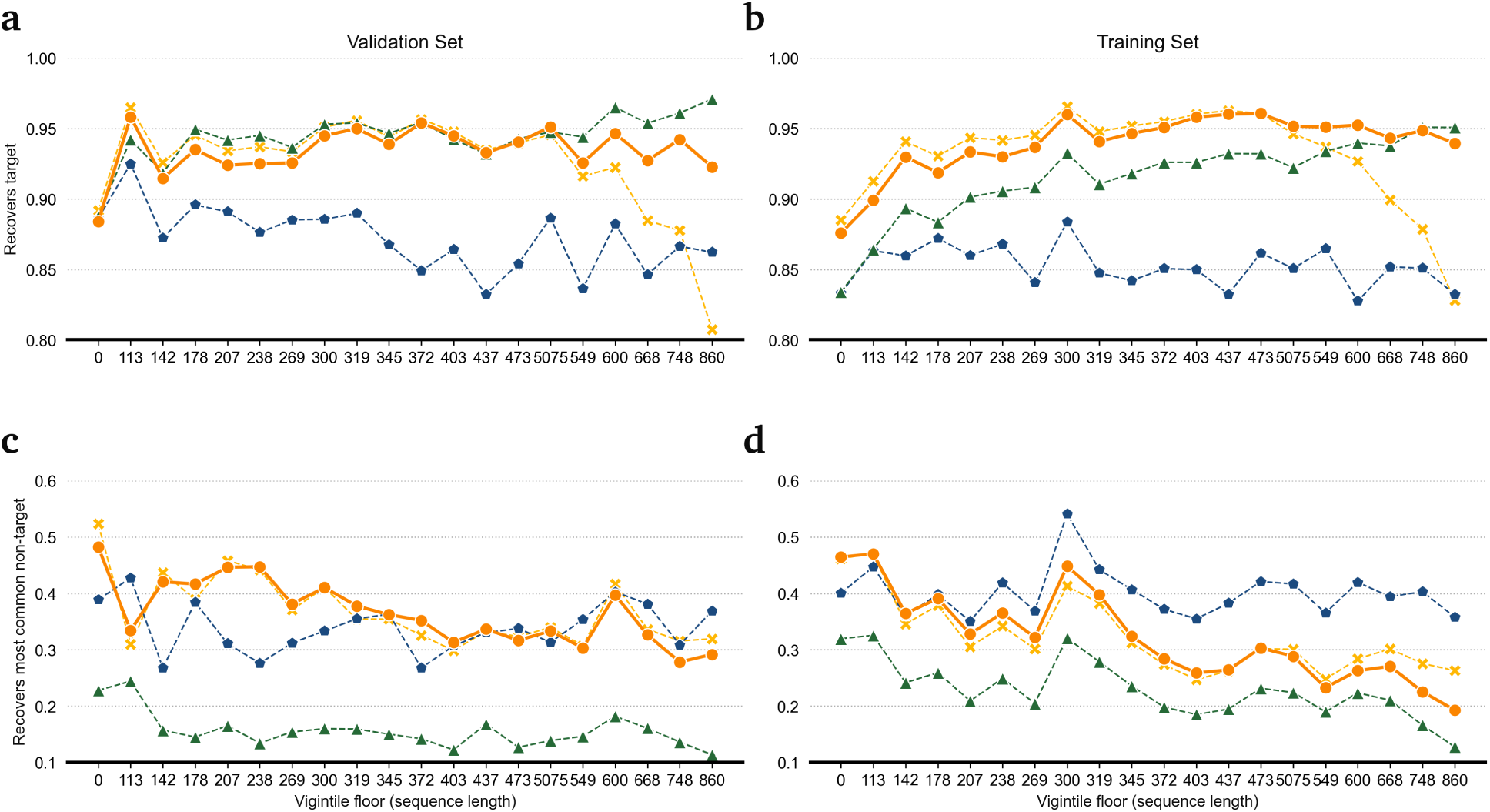
Context extension. The lines represent sequence recovery statistics for Category B residues at the top (’Recovers target’) and Category I residues at the bottom (’Recovers most common non-target’) across four models. ESM2-15B is depicted by the dashed blue line with pentagon symbols, while ProGen2-xlarge is shown by the dashed green line with triangles. The base 350M parameter AMPLIFY model (before context extension) is indicated by the dashed yellow line with crosses, and the extended context AMPLIFY 350M model is shown as a solid orange line with circles. Data points represent recovery statistics over an equivalent number of sequences, with proteins divided into 20 evenly sized sets (vigintiles) based on sequence length.

Our tests for recovering the most common non-target residues in Categories E-I measure the model’s ability to predict synonyms that may be more desirable than the target residue. This goal is inherently at odds with the training task, and across all model families, we observe that improving target sequence recovery by increasing model scale comes at a cost. Overtraining on mask replacement reduces the model’s ability to predict non-target residues that we expect to be not just valid but superior synonyms.

For AMPLIFY, we aimed to optimize sequence recovery by adjusting the data and only scaled up to the point where Category B accuracy was saturated. This approach is significantly more efficient, both in training and deployment, and helps avoid the overfitting associated with scaling up.

When considering both tasks as valid goals and comparing the sum recovery of either (Figures 3 and 4, bottom panels), the closest comparable model to AMPLIFY 350M is ProGen2-xlarge, a decoder-style model that is 18.5× larger. While these two models perform similarly across all categories when considering both tasks together, ProGen2 achieves this by prioritizing target sequence recovery. We argue that this is not advantageous, as it indicates overtraining on the wrong task.

#### Training Data Scale

One of the initial steps in our training program was to establish controls by retraining ESM2 models using our data and test sets. We found that simply updating the data led to a significant improvement in model performance. As discussed in the main text, this reflects the ongoing community-driven enhancement of protein databases.

We observe a significant dip in model quality when training on UniRef versions released between 2018 and 2021, which we attribute to specific sequences that have since been removed from the data, as discussed in the main text. This process of sequence culling over time is a notable characteristic of the datasets; for example, 31% of the UniProt sequences in UniRef 2015 have been specifically annotated as deleted accessions in the years since.

However, looking beyond sequence curation, we find that performance on target sequence recovery has not changed significantly in recent years compared to 2011-2015 (Figure 5, Top). What has changed is performance on our orthogonal task (Figure 5, Middle), where we see a consistent decline in performance as new data is added for test set sequences, and a consistent improvement for training set sequences.

This indicates that the primary benefit of adding additional sequences is to mitigate overtraining, particularly overtraining on specific sequences within the training set. While this is expected behavior, it is not fully captured by the loss function we optimize during training.

The expectation that larger training data sets will prevent overfitting largely stems from the fact that protein sequences are not independent of one another. Protein sequence data is generated almost entirely through error-prone inheritance and selection for fitness, representing hundreds of millions of years of sequential modification and direct relationships. As a result, when our data expands, it primarily grows by adding functional synonyms. Learning these relationships seems to be at odds with our current training task.

### Context Extension

During the first 1 million training steps, we use a context window of 512 because most protein sequences in our training data fit within this window, increasing efficiency by avoiding unnecessary padding. For proteins longer than this, sequences are randomly trimmed to fit, which ensures that they are seen during training. However we did notice a performance drop on longer proteins. We hypothesized that this issue partly stemmed from the model’s difficulty in correctly extrapolating the RoPE positional embeddings to longer contexts. To address this, we extended training by 25k steps for the 120M model and 50k steps for the 350M model using a context length of 2048. This adjustment successfully resolved the problem.

Figure 6 illustrates this for the 350M parameter model, comparing its performance on specific protein lengths to comparable models (ESM2-15B and ProGen2-xlarge). The plots also indicate that the performance issue is specific to mask replacement, as the ability to generalize toward predicting the most common non-target identities remains unaffected. This suggests that we are primarily correcting a memorization issue.

Among these models, the trade-off between target sequence recovery and most-common non-target recovery is evident. ProGen2-xlarge shows a clear advantage in the first task for long proteins within our test set but has the worst performance in the second task. Conversely, ESM2 generally performs worse in the first task but excels in the second task for proteins within AMPLIFY’s training data.

It’s important to note that the AMPLIFY test sets are excluded only when training AMPLIFY. The comparisons with ProGen2 and ESM2 show performance on the same sets but do not account for the effect of excluding sequences from training. We do not have access to the specific sequences ProGen2 and ESM2 were trained on, and in the case of ESM2, the performance on non-target recovery might reflect differences in exclusion due to the use of clustered sequences.

### Embedding Analysis

#### Emergent Non-Protein Prediction

In exploring model behavior, we tested a variety of statistical methods, dimensionality reduction, similarity clustering, and visualization. Matching expectations from prior protein LLMs, the primary observation when looking at the residue embedding values (taken from the last layer of network weights) is that positions the model is confident in separate into 20 broadly distributed clusters corresponding to the 20 amino acid types, while low confidence predictions instead show higher similarity to one another in the embedding space, forming blended clusters that often appear as extensions of the confident position clusters.

To illustrate this effect, we chose to isolate data for a single residue type within a sequence motif of known function, specifically the arginine residue of RG motifs, which is a well-characterized RNA binding motif that is found and functional in both folded and intrinsically disordered proteins. We identify 36,580 of these positions in the subset of the human proteome with AlphaFold2 structure predictions and in Figures 2A-C and Figures 7A-C show UMAP projections demonstrating that the embedding space contains a tightly clustered extension of residues with low confidence AlphaFold2 predictions, no gene ontology annotation data for molecular localization, and no direct evidence of protein existence.

**Figure 7:**
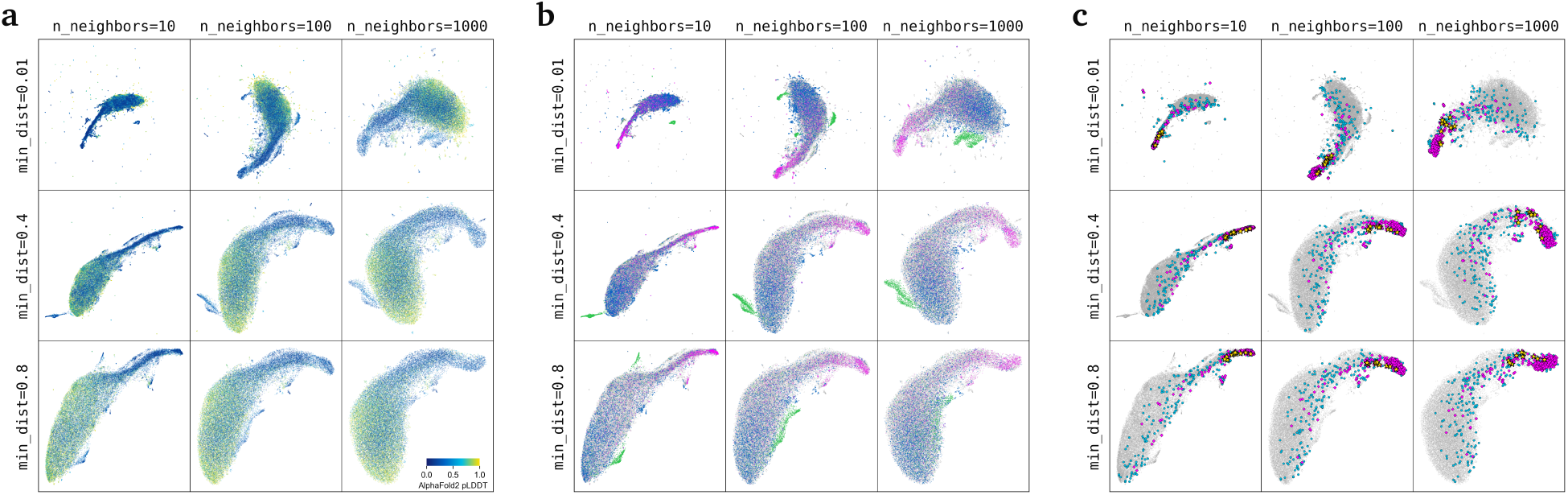
Non-protein extension is independent of parameter choice. Panels a-c show UMAP parameter effects on the RG motif projections shown in main text figure 2a-c, with the main text parameters of min_dist=0.4 and n_neighbors=100 in the center. Projections all show a tightly clustered set of residues separate from the bulk of the data, with this set showing a) low confidence predictions by AlphaFold2, b) no annotation data in the gene ontology database, and c) localization to the same region as non-protein sequence data (yellow stars showing RG motifs from the text of Frankenstein).

To serve as a non-protein sequence control we established a conversion function that strips out characters not found in protein sequences and used that to encode sentences from human literature, with results shown here for the first chapter of Frankenstein, the novel by Mary Shelley. These non-protein controls show that the embedding space occupied by many un-annotated and hypothetical proteins is also occupied when embedding sequences that are not proteins at all and are not part of the training corpus.

While it goes without saying that a language model trained on proteins does not display any ability to perform mask replacement tasks on natural language text, it is worth looking at how a protein language model embeds it. The embeddings for different tokens in non-protein text show very high similarity to one another, even when unmasked, and appear to converge on a specific region of the landscape, potentially representing a catch-all null region for tokens that the model cannot identify any information on.

This feature of model behavior gives us two different ways to establish that a given full protein sequence doesn’t fit on the known protein space learned by the model, as shown in Figure 2D. Hypothetical proteins are significantly more likely to show two related traits, one of them is similarity to non-human text, but the other is just convergence. If you compare similarity between each residue embedding in a given sequence, the proteins that show similarity to human text are consistently also the ones where every residue embedding converge on a tightly defined mean.

We hypothesize that one of the things the model learns is when to ignore sequence context because the sequence is fundamentally unpredictable, either because it’s too distant from anything else in the training data or because it doesn’t follow the basic rules governing protein sequence evolution. Manual inspection shows that many of the human reference proteome sequences that fall in this space can be confirmed as non-coding, and that a significant fraction of our training data may need to be reevaluated. Identifying what the empirical evidence is behind each sequence identified in this way provides a clear path to future development.

### Comparison to Structural Models

#### Recognizing Disordered Proteins

Our previous example on RG motifs (Figures 7A-C) shows that low confidence AlphaFold2 predictions occur for residues spanning a wide range of protein space. For example, AlphaFold2 does not confidently predict the structure of collagen chains, and that can be observed in the pLM embedding space as a distinct cluster separate from non-proteins.

Low confidence predictions have been described in the literature as a way to identify disordered proteins, but it is more accurately just a description of sequences that the model doesn’t recognize, which given the fact the AlphaFold2 trains on structured proteins includes both disordered proteins and non-proteins. That distinction is very important because disordered proteins aren’t just random sequences, empirical testing shows that the majority of sequence space is insoluble, so natural proteins that can be both soluble and disordered are a specific evolved class. Using AlphaFold2 confidence values to predict disorder conflates these two distinct classes into one.

To demonstrate this, we created a negative set of 296 proteins from the human reference proteome that are at high risk of being non-proteins, with protein evidence level 5 and no annotation data in the gene ontology database for cellular localization and a positive set of 240 human proteins with < 25% of their residues annotated in the DisProt database as being intrinsically disordered. Using ROC-AUC analysis (Figure 2H) we observe that AlphaFold2 confidence has no ability to discriminate these two populations from one another, but a simple test of pLM similarity to non-protein text can easily separate the two.

### Structural Feature Prediction

To test discrimination between structural features, we prepared a zero-shot task by using the residue embedding of human proteins to identify training set residues that are similar to test set residues. Using the AlphaFold2 structures of high confidence human proteins (PE=1) and a method where cosine similarity of AMPLIFY 350M embeddings for all applicable test set residues (N=849,055) is calculated against all applicable training set residues (N=4,196,541) we can take the most similar hit (KNN with a K of 1) from a completely different human protein and assess how predictive the properties of the training set residues are for the test set residues.

We demonstrate that 3-state DSSP [Kabsch and Sander, 1983, Touw et al., 2014] secondary assignments (helix, strand, and loop) can be predicted using this method at an accuracy that ranges from 66-91% depending on the confidence value (pLDDT) AlphaFold2 provides for its predictions (Figure 8A). This range, in part, reflects the secondary structure distribution changes seen for residues predicted at different confidence regimes. This appears to be because low pLDDT residues are usually assigned to loop conformations by DSSP, and we observe that this zero shot predictor is also adept at predicting which residues will show up at lower pLDDT (Figure 8B), with the nearest neighbor’s pLDDT value predicting the target’s at an AUC of 0.93 for high confidence (pLDDT < 90) vs low confidence (pLDDT < 60).

**Figure 8:**
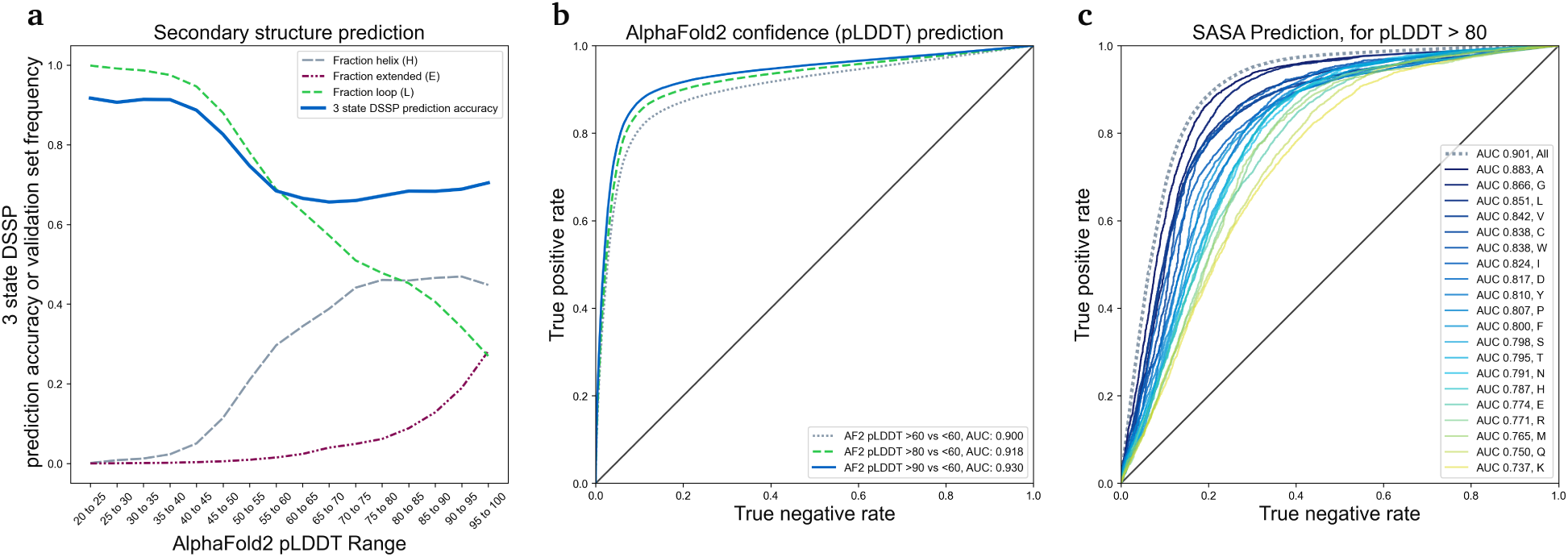
Structural property prediction. KNN prediction values for human test set residues with AlphaFold2 structures and protein evidence level 1. Panel a) shows the frequency of three state DSSP assignments for residues divided into bins by their pLDDT prediction confidence values, with dashed lines showing a conversion from primarily loop character (green) to helical character (gray) with some beta strand (burgundy) as pLDDT increases. The solid blue line then shows KNN prediction accuracy for the three-class labels observed in each pLDDT bin. Panel b) shows ROC curves for discriminating between high-confidence and low-confidence AlphaFold2 predictions, where more stringent confidence thresholds result in higher predictivity. With negative class is pLDDT < 60, lines show positive class < 60 (gray dots), < 80 (green dashes), and < 90 (solid blue). Panel c) shows ROC analysis for discriminating between the top 25% (positive set) and bottom 25% (negative set) of residues by SASA. The dashed gray line shows performance over all residues, and solid-colored lines show the same analysis done for each residue type independently, with lines ordered from blue to yellow by descending ROC-AUC performance.

**Figure 9:**
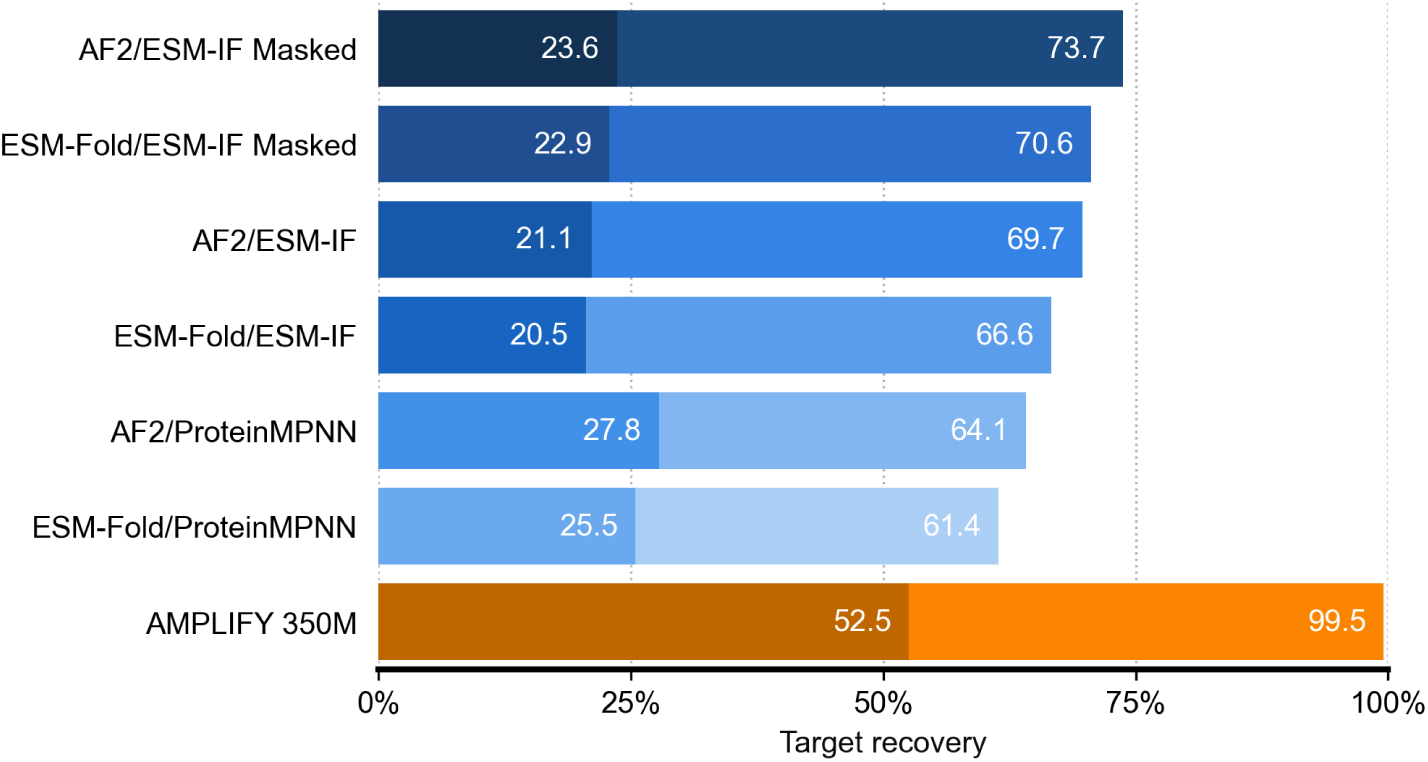
Sequence-to-structure-to-sequence prediction. Sequence recovery accuracy on human proteome binned by sequence conservation. Light bars show recovery of the target sequence at positions with 96-99% sequence conservation, and shaded bars show recovery of the most common non-target sequence when found at > 70% conservation. Protocols involving both a structure prediction model and a structure-to-sequence model are labeled by both methods divided by a slash.

For the highest confidence predictions, the 3-state secondary structure prediction only hits 70% accuracy, which isn’t the state of the art that can be obtained when actually training a predictor. As a zero-shot test, what this demonstrates is that even without training on this task, the embedded landscape is placing residues that are found in helices closer to other residues found in helices, a phenomenon suggestive of emergent learning.

Other properties are also predictable in this way. For solvent-accessible surface area (calculated by Biotite [Shrake and Rupley, 1973, Kunzmann and Hamacher, 2018]), taken only for high confidence predictions (pLDDT > 90), we can separate the most accessible (top 25% by SASA) residues from the most buried (bottom 25% by SASA) at a ROC-AUC of 0.90. That value though partially reflects high accuracy for predicting the protein sequence itself (which at 93% means most of the residues are matched correctly), but if you split out the 20 amino acids to separate measurements, all 20 are independently predictive, with AUC values ranging from 0.74 to 0.88 (Figure 8C).

The nature of vector space lookup prediction introduces new possibilities worth further exploration. While the task itself requires a significant upfront cost to establish a database of vectors, the similarity lookup task itself very fast, the computational step of calculating cosine similarity for one residue to all 84 million residues in our KNN set takes less than 0.01 seconds (when done as a single PyTorch matrix operation on an A100 80GB) and significant optimization is possible with a vector database backend. The task itself does not require the generation of a multiple sequence alignment and, by its nature, can produce fully interpretable predictions.

### Utility in Protein Design

A common workflow in computational protein design involves using protein structure predictors like AlphaFold2 [Jumper et al., 2021] and ESM-Fold [Lin et al., 2023] to generate structures for designed sequences, followed by using structure-to-sequence models, such as ProteinMPNN [Dauparas et al., 2022] or ESM-IF [Hsu et al., 2022], to generate sequences compatible with those structures. Newer models, like AlphaFold3 [Abramson et al., 2024] and ESM3 [Hayes et al., 2024], claim to perform both tasks simultaneously, but their licensing restrictions make them difficult to test. However, we can evaluate the existing workflows.

For our human proteome test proteins, we created two sets of structures: an ESM-Fold set generated by running the model on available sequences using default parameters, and an AlphaFold2 set downloaded directly from the UniProt AlphaFold database. During testing, we discovered that about 2% of the AlphaFold downloads do not match the sequences currently associated with their UniProt accessions, as the sequences have been updated since the structures were initially deposited. Rather than excluding these sequences, we replaced the missing structures in this set with their ESM-Fold counterparts (these proteins are annotated as ‘AF2 replaced’ in the supplementary data files).

We then established three methods for predicting masked sequences using different structure-to-sequence models. For two of these methods, we used ProteinMPNN and ESM-IF with their default settings. Under default conditions these models use fully masked sequences, relying solely on backbone coordinate information from the structure as input, although sequence information is part of the input context that is usually derived entirely from the structure. To create a method more directly comparable to our pLM testing approach, we developed a third method (ESM-IF Masked), where ESM-IF is provided with both the structure and the preceding sequence when predicting logits for each position.

These tests lead to three main conclusions. First, despite being an incomplete set, AF2 structures consistently outperform in recovering both target sequences and the most common non-target sequences. Second, ESM-IF surpasses ProteinMPNN in target sequence recovery but falls short in recovering the most common non-target sequences. Third, while providing full decoder-style sequence context (as in ESM-IF Masked) does improve performance, it still doesn’t make this approach competitive with pLM performance, as exemplified by AMPLIFY 350M.

While these methods have proven valuable in protein design, they may not be as effective across the diverse range of proteins found in the human proteome. ProteinMPNN and ESM-IF are trained on protein structures and typically tested on the design of individual, well-folded domains. The differences in recovery might reflect variations in model performance, but they could also indicate how limited these structure-based tasks are in capturing the full complexity of human protein space.

### Unsupervised Contact Prediction

Previous studies [Rao et al., 2020, Lin et al., 2023] demonstrated that attention maps in bidirectional Transformer pLMs capture structural information, enabling the linear prediction of protein contacts with minimal supervision for calibration. This section shows that structural information also emerges within AMPLIFY attention maps.

In order to ensure a fair comparison, we closely followed the methodology of previous works. Specifically, for a protein with n residues and a pLM with L layers and K attention heads per layer, we extract the resulting LK attention maps, denoted as 𝑨_𝑙𝑘_ = Softmax(𝑸𝑲^⊤^), each of size (𝑛, 𝑛). These attention maps are independently symmetrized and corrected using the average product correction (APC) defined as follows:

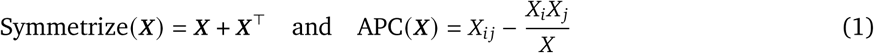

where 𝑋_𝑖_, 𝑋_𝑗_, and 𝑿 represent the sums over the 𝑖-th row, 𝑗-th column, and the entire matrix, respectively. A logistic regression is fitted on the processed attention maps of a few proteins for calibration purposes. Note that one coefficient is learned per head.

As in prior works, a contact is defined as a pair of amino acids with a C-alpha distance of less than 8Å. The logistic regression is regularized with an L1 penalty (𝜆 = 0.15), and a random subset of 20 proteins from each set is selected to train the classifier. This process excludes local contacts by considering amino acid pairs where |𝑖 − 𝑗| < 6. Predictions are evaluated using the long-range precision at L (LR P@L). Given a protein with L amino acids, the LR P@L metric is defined as the precision of the top L predicted contacts where |𝑖 − 𝑗| ≥ 24. The evaluation is done with the ESM codebase for consistency.

Table 8 shows that our replication of the methodology achieves comparable LR P@L on CASP14 to that of Lin et al. [2023]. While they evaluated the Apr-Jun 2022 release of CAMEO, AMPLIFY was trained on data up to December 2023. In order to prevent data leakage and thus overestimate the performance, we evaluated all models on the most recent CAMEO release available at the time of writing. Additionally, we included results from the more recent CASP15 dataset.

Interestingly, while AMPLIFY outperforms even the best ESM on sequence recovery, ESM outperforms AMPLIFY at the task of unsupervised contact prediction. Our investigation into potential factors allowed us to rule out suboptimal hyperparameters, the lack of calibration samples, and FlashAttention instability [Golden et al., 2024]. A key distinction of AMPLIFY’s architecture compared to ESM2 is its smaller number of attention heads. As ESM’s methodology relies on these heads for contact prediction, AMPLIFY may be at a disadvantage. As a result, structural information might not emerge as clearly in the attention maps in a linear fashion. In order to explore this hypothesis, we replace the logistic regression with a random forest. Note that the calibration remained limited to the same 20 proteins as for the logistic regression.

Table 9 reveals that the non-linear random forest improves the performance of all pLMs, with the greatest improvement observed in AMPLIFY. On CASP14, random forest improves the LR P@L of ESM2-15B by 38% and AMPLIFY 350M by 146% compared to the logistic regression. However, AMPLIFY still lags behind ESM2 at similar number of parameters, indicating that the non-linearity of the information is not the main reason for the discrepency. To further explore this hypothesis, we perform contact prediction with the categorical Jacobian proposed by Zhang et al. [2024], which quantifies of all possible mutations on all positions. Given the high computational cost of this method, ESM2 15B was excluded from this analysis. Additionally, we consider medium-and short-range interactions and report the AUC in addition to the P@L.

Table 10 suggests that the number of attention heads is not the main limiting factor. Instead, we hypothesise that overfitting, in the sense of deviating from consensus, helps at contact prediction because the model needs to pay attention to specific patterns both in the attention maps and input-output. While AMPLIFY is not competitive with ESM2 at directly predicting contacts, many tasks including protein design benefit from aligning with the consensus.

**Table 10:** Contact prediction using the Categorical Jacobian. The standard deviation is computed across the proteins in each dataset.

|  |  | Short |  | Medium |  | Long |  |
| --- | --- | --- | --- | --- | --- | --- | --- |
|  |  | (6 < \|i - j\| < 12) |  | (12 < \|i - j\| < 24) |  | (24 < \|i - j\|) |  |
|  |  | AUC | P@L | AUC | P@L | AUC | P@L |
| CASP14 | ESM2 8M | 7.6 ± 4.9 | 7.3 ± 4.7 | 10.6 ± 9.2 | 8.5 ± 6.4 | 8.5 ± 10.7 | 7.0 ± 8.1 |
|  | ESM2 35M | 11.2 ± 7.5 | 9.0 ± 6.1 | 12.3 ± 10.4 | 9.6 ± 7.6 | 11.6 ± 12.6 | 9.1 ± 9.6 |
|  | ESM2 150M | 12.2 ± 10.6 | 10.0 ± 6.9 | 17.3 ± 14.7 | 12.9 ± 10.0 | 18.0 ± 15.1 | 13.9 ± 12.1 |
|  | ESM2 650M | 20.0 ± 15.2 | 14.3 ± 9.9 | 26.3 ± 18.1 | 18.3 ± 12.0 | 28.7 ± 22.5 | 23.0 ± 18.4 |
|  | ESM2 3B | 19.3 ± 13.1 | 15.4 ± 9.9 | 26.5 ± 17.5 | 18.9 ± 12.0 | 29.0 ± 20.7 | 22.8 ± 16.2 |
|  | AMPLIFY 120M | 9.6 ± 6.6 | 8.9 ± 5.7 | 12.2 ± 10.9 | 9.8 ± 7.7 | 9.7 ± 11.9 | 7.8 ± 9.0 |
|  | AMPLIFY 350M | 11.3 ± 8.0 | 9.2 ± 5.8 | 13.8 ± 11.9 | 10.5 ± 8.8 | 10.8 ± 11.8 | 9.1 ± 8.9 |
| CASP15 | ESM2 8M | 6.9 ± 6.3 | 6.3 ± 4.3 | 11.0 ± 10.7 | 8.8 ± 7.8 | 6.3 ± 8.7 | 5.4 ± 5.9 |
|  | ESM2 35M | 12.3 ± 8.5 | 9.4 ± 5.2 | 17.6 ± 14.8 | 13.2 ± 10.6 | 20.5 ± 14.4 | 15.9 ± 10.6 |
|  | ESM2 150M | 17.0 ± 9.5 | 12.2 ± 5.3 | 23.5 ± 17.3 | 16.4 ± 11.3 | 24.9 ± 14.5 | 19.7 ± 10.7 |
|  | ESM2 650M | 20.6 ± 10.9 | 14.1 ± 5.8 | 26.2 ± 17.5 | 18.4 ± 11.5 | 24.3 ± 15.6 | 20.9 ± 12.6 |
|  | ESM2 3B | 20.6 ± 10.6 | 13.9 ± 5.6 | 24.6 ± 16.6 | 17.5 ± 10.3 | 24.4 ± 16.1 | 21.4 ± 12.3 |
|  | AMPLIFY 120M | 10.1 ± 5.7 | 8.0 ± 3.7 | 12.7 ± 12.3 | 9.5 ± 8.7 | 12.4 ± 13.4 | 9.9 ± 10.5 |
|  | AMPLIFY 350M | 11.4 ± 8.0 | 9.1 ± 4.4 | 13.5 ± 13.1 | 9.6 ± 8.1 | 14.2 ± 12.9 | 10.8 ± 9.8 |
| CAMEO | ESM2 8M | 7.1 ± 6.9 | 6.7 ± 5.1 | 6.9 ± 8.1 | 8.5 ± 10.8 | 12.7 ± 14.6 | 9.8 ± 10.5 |
|  | ESM2 35M | 10.1 ± 9.0 | 8.5 ± 6.4 | 13.2 ± 13.3 | 10.1 ± 9.9 | 18.5 ± 17.7 | 14.3 ± 13.1 |
|  | ESM2 150M | 14.7 ± 10.5 | 11.3 ± 7.3 | 19.4 ± 15.5 | 14.6 ± 12.0 | 23.4 ± 18.0 | 19.0 ± 14.0 |
|  | ESM2 650M | 17.4 ± 10.6 | 12.6 ± 7.2 | 21.1 ± 15.6 | 16.2 ± 12.2 | 23.8 ± 17.0 | 20.4 ± 13.6 |
|  | ESM2 3B | 17.7 ± 10.2 | 12.8 ± 6.6 | 21.0 ± 15.1 | 15.8 ± 11.3 | 24.2 ± 16.2 | 20.7 ± 13.1 |
|  | AMPLIFY 120M | 9.7 ± 8.5 | 8.4 ± 6.0 | 12.3 ± 11.7 | 9.7 ± 8.9 | 15.7 ± 14.3 | 12.5 ± 10.8 |
|  | AMPLIFY 350M | 10.6 ± 8.1 | 9.1 ± 5.9 | 13.6 ± 11.3 | 10.6 ± 8.9 | 16.7 ± 14.0 | 13.5 ± 11.0 |

## Footnotes

1 The inclusion of the paired OAS sequences and SCOP domains increases the size of UR100 by <1%.

2 A larger model with strong regularization or fewer training steps would also reduce memorization, but would be more costly at inference and produce large embeddings unsuitable for downstream tasks with limited data.

3 https://doi.org/10.5281/zenodo.13834052

4 https://huggingface.co/chandar-lab

5 https://github.com/chandar-lab/AMPLIFY

6 https://huggingface.co/chandar-lab

7 https://www.nvidia.com/content/dam/en-zz/Solutions/Data-Center/a100/pdf/a100-80gb-datasheet-update-nvidia-us-1521051-r2-web.pdf

8 https://github.com/facebookresearch/esm/discussions/414

9 https://aws.amazon.com/ec2/instance-types/p4/

10 https://aws.amazon.com/ec2/instance-types/p3/

11 https://images.nvidia.com/aem-dam/en-zz/Solutions/data-center/nvidia-ampere-architecture-whitepaper.pdf

12 Accessions that have been deleted as of the 2024_3 release of UniProt (May 29, 2024) indicate that either their source DNA sequence was removed or they were reclassified as incorrectly predicted to code for a protein.

## References

Josh Abramson, Jonas Adler, Jack Dunger, Richard Evans, Tim Green, Alexander Pritzel, Olaf Ronneberger, Lindsay Willmore, Andrew J Ballard, Joshua Bambrick, et al. Accurate structure prediction of biomolecular interactions with alphafold 3. Nature, pages 1–3, 2024.

Reza Yazdani Aminabadi, Samyam Rajbhandari, Ammar Ahmad Awan, Cheng Li, Du Li, Elton Zheng, Olatunji Ruwase, Shaden Smith, Minjia Zhang, Jeff Rasley, et al. Deepspeed-inference: enabling efficient inference of transformer models at unprecedented scale. In SC22: International Conference for High Performance Computing, Networking, Storage and Analysis, pages 1–15. IEEE, 2022.

Christian B Anfinsen. Principles that govern the folding of protein chains. Science, 181(4096):223–230, 1973.

Frances H Arnold. Directed evolution: bringing new chemistry to life. Angewandte Chemie (International Ed. in English), 57(16):4143, 2018.

Michael Ashburner, Catherine A Ball, Judith A Blake, David Botstein, Heather Butler, J Michael Cherry, Allan P Davis, Kara Dolinski, Selina S Dwight, Janan T Eppig, et al. Gene ontology: tool for the unification of biology. Nature genetics, 25(1):25–29, 2000.

Maria Cristina Aspromonte, Maria Victoria Nugnes, Federica Quaglia, Adel Bouharoua, Silvio CE Tosatto, and Damiano Piovesan. Disprot in 2024: improving function annotation of intrinsically disordered proteins. Nucleic Acids Research, 52(D1):D434–D441, 2024.

Salman F Banani, Hyun O Lee, Anthony A Hyman, and Michael K Rosen. Biomolecular condensates: organizers of cellular biochemistry. Nature reviews Molecular cell biology, 18(5):285–298, 2017.

Patrick Bryant, Gabriele Pozzati, and Arne Elofsson. Improved prediction of protein-protein interactions using alphafold2. Nature communications, 13(1):1265, 2022.

Nicholas Carlini, Daphne Ippolito, Matthew Jagielski, Katherine Lee, Florian Tramer, and Chiyuan Zhang. Quantifying memorization across neural language models. In The Eleventh International Conference on Learning Representations, 2023. URL https://openreview.net/forum?id=TatRHT_1cK.

John-Marc Chandonia, Lindsey Guan, Shiangyi Lin, Changhua Yu, Naomi K Fox, and Steven E Brenner. Scope: improvements to the structural classification of proteins–extended database to facilitate variant interpretation and machine learning. Nucleic acids research, 50(D1):D553–D559, 2022.

The UniProt Consortium. Uniprot: the universal protein knowledgebase in 2023. Nucleic Acids Research, 51 (D1):D523–D531, 11 2022. ISSN 0305-1048. doi: 10.1093/nar/gkac1052.

Tri Dao, Dan Fu, Stefano Ermon, Atri Rudra, and Christopher Ré. Flashattention: Fast and memory-efficient exact attention with io-awareness. Advances in Neural Information Processing Systems, 35:16344–16359, 2022.

J. Dauparas, I. Anishchenko, N. Bennett, H. Bai, R. J. Ragotte, L. F. Milles, B. I. M. Wicky, A. Courbet, R. J. de Haas, N. Bethel, P. J. Y. Leung, T. F. Huddy, S. Pellock, D. Tischer, F. Chan, B. Koepnick, H. Nguyen, A. Kang, B. Sankaran, A. K. Bera, N. P. King, and D. Baker. Robust deep learning–based protein sequence design using proteinmpnn. Science, 378(6615):49–56, 2022. doi: 10.1126/science.add2187. URL https://www.science.org/doi/abs/10.1126/science.add2187.

Jacob Devlin, Ming-Wei Chang, Kenton Lee, and Kristina Toutanova. BERT: Pre-training of deep bidirectional transformers for language understanding. In Jill Burstein, Christy Doran, and Thamar Solorio, editors, Proceedings of the 2019 Conference of the North American Chapter of the Association for Computational Linguistics: Human Language Technologies, Volume 1 (Long and Short Papers), pages 4171–4186, Minneapolis, Minnesota, June 2019. Association for Computational Linguistics. doi: 10.18653/v1/N19-1423. URL https://aclanthology.org/N19-1423.

Dimiter S Dimitrov. Therapeutic proteins. Therapeutic Proteins: Methods and Protocols, pages 1–26, 2012.

Sean R. Eddy. Profile hidden markov models. Bioinformatics (Oxford, England), 14(9):755–763, 1998.

Noelia Ferruz and Birte Höcker. Controllable protein design with language models. Nature Machine Intelligence, 4(6):521–532, 2022.

Leo Gao, Stella Biderman, Sid Black, Laurence Golding, Travis Hoppe, Charles Foster, Jason Phang, Horace He, Anish Thite, Noa Nabeshima, et al. The pile: An 800gb dataset of diverse text for language modeling. arXiv preprint arXiv:2101.00027, 2020.

Alicia Golden, Samuel Hsia, Fei Sun, Bilge Acun, Basil Hosmer, Yejin Lee, Zachary DeVito, Jeff Johnson, Gu-Yeon Wei, David Brooks, et al. Is flash attention stable? arXiv preprint arXiv:2405.02803, 2024.

Thomas Hayes, Roshan Rao, Halil Akin, Nicholas J. Sofroniew, Deniz Oktay, Zeming Lin, Robert Verkuil, Vincent Q. Tran, Jonathan Deaton, Marius Wiggert, Rohil Badkundri, Irhum Shafkat, Jun Gong, Alexander Derry, Raul S. Molina, Neil Thomas, Yousuf Khan, Chetan Mishra, Carolyn Kim, Liam J. Bartie, Matthew Nemeth, Patrick D. Hsu, Tom Sercu, Salvatore Candido, and Alexander Rives. Simulating 500 million years of evolution with a language model. bioRxiv, 2024. doi: 10.1101/2024.07.01.600583. URL https://www.biorxiv.org/content/early/2024/07/02/2024.07.01.600583.

Jordan Hoffmann, Sebastian Borgeaud, Arthur Mensch, Elena Buchatskaya, Trevor Cai, Eliza Rutherford, Diego de Las Casas, Lisa Anne Hendricks, Johannes Welbl, Aidan Clark, et al. Training compute-optimal large language models. arXiv preprint arXiv:2203.15556, 2022.

Chloe Hsu, Robert Verkuil, Jason Liu, Zeming Lin, Brian Hie, Tom Sercu, Adam Lerer, and Alexander Rives. Learning inverse folding from millions of predicted structures. In Kamalika Chaudhuri, Stefanie Jegelka, Le Song, Csaba Szepesvari, Gang Niu, and Sivan Sabato, editors, Proceedings of the 39th International Conference on Machine Learning, volume 162 of *Proceedings of Machine Learning Research*, pages 8946–8970. PMLR, 17–23 Jul 2022. URL https://proceedings.mlr.press/v162/hsu22a.html.

John B Ingraham, Max Baranov, Zak Costello, Karl W Barber, Wujie Wang, Ahmed Ismail, Vincent Frappier, Dana M Lord, Christopher Ng-Thow-Hing, Erik R Van Vlack, et al. Illuminating protein space with a programmable generative model. Nature, 623(7989):1070–1078, 2023.

John Jumper, Richard Evans, Alexander Pritzel, Tim Green, Michael Figurnov, Olaf Ronneberger, Kathryn Tunyasuvunakool, Russ Bates, Augustin Žídek, Anna Potapenko, et al. Highly accurate protein structure prediction with alphafold. nature, 596(7873):583–589, 2021.

Wolfgang Kabsch and Christian Sander. Dictionary of protein secondary structure: Pattern recognition of hydrogen-bonded and geometrical features. Biopolymers, 22(12):2577–2637, 1983. doi: 10.1002/bip.360221211. URL https://onlinelibrary.wiley.com/doi/abs/10.1002/bip.360221211.

Hetunandan Kamisetty, Arvind Ramanathan, Chris Bailey-Kellogg, and Christopher James Langmead. Account-ing for conformational entropy in predicting binding free energies of protein-protein interactions. Proteins: Structure, Function, and Bioinformatics, 79(2):444–462, 2011.

Jared Kaplan, Sam McCandlish, Tom Henighan, Tom B Brown, Benjamin Chess, Rewon Child, Scott Gray, Alec Radford, Jeffrey Wu, and Dario Amodei. Scaling laws for neural language models. arXiv preprint arXiv:2001.08361, 2020.

Aleksandr Kovaltsuk, Jinwoo Leem, Sebastian Kelm, James Snowden, Charlotte M Deane, and Konrad Krawczyk. Observed antibody space: a resource for data mining next-generation sequencing of antibody repertoires. The Journal of Immunology, 201(8):2502–2509, 2018.

Patrick Kunzmann and Kay Hamacher. Biotite: A unifying open source computational biology framework in python. BMC Bioinformatics, 19, 10 2018. doi: 10.1186/s12859-018-2367-z.

Denis LJ Lafontaine, Joshua A Riback, Rümeyza Bascetin, and Clifford P Brangwynne. The nucleolus as a multiphase liquid condensate. Nature reviews Molecular cell biology, 22(3):165–182, 2021.

Martin Lehmann, L Pasamontes, SF Lassen, and M Wyss. The consensus concept for thermostability engineering of proteins. Biochimica et Biophysica Acta (BBA)-protein structure and molecular enzymology, 1543(2): 408–415, 2000.

Yoav Levine, Noam Wies, Or Sharir, Hofit Bata, and Amnon Shashua. Limits to depth efficiencies of self-attention. Advances in Neural Information Processing Systems, 33:22640–22651, 2020.

Zeming Lin, Halil Akin, Roshan Rao, Brian Hie, Zhongkai Zhu, Wenting Lu, Allan dos Santos Costa, Maryam Fazel-Zarandi, Tom Sercu, Sal Candido, et al. Language models of protein sequences at the scale of evolution enable accurate structure prediction. BioRxiv, 2022:500902, 2022.

Zeming Lin, Halil Akin, Roshan Rao, Brian Hie, Zhongkai Zhu, Wenting Lu, Nikita Smetanin, Robert Verkuil, Ori Kabeli, Yaniv Shmueli, et al. Evolutionary-scale prediction of atomic-level protein structure with a language model. Science, 379(6637):1123–1130, 2023.

I Loshchilov. Decoupled weight decay regularization. arXiv preprint arXiv:1711.05101, 2017.

John S Mattick, Paulo P Amaral, Piero Carninci, Susan Carpenter, Howard Y Chang, Ling-Ling Chen, Runsheng Chen, Caroline Dean, Marcel E Dinger, Katherine A Fitzgerald, et al. Long non-coding rnas: definitions, functions, challenges and recommendations. Nature reviews Molecular cell biology, 24(6):430–447, 2023.

Paulius Micikevicius, Sharan Narang, Jonah Alben, Gregory Diamos, Erich Elsen, David Garcia, Boris Ginsburg, Michael Houston, Oleksii Kuchaiev, Ganesh Venkatesh, et al. Mixed precision training. arXiv preprint arXiv:1710.03740, 2017.

Tanja Mittag, Lewis E Kay, and Julie D Forman-Kay. Protein dynamics and conformational disorder in molecular recognition. Journal of Molecular Recognition: An Interdisciplinary Journal, 23(2):105–116, 2010.

John Moult, Krzysztof Fidelis, Andriy Kryshtafovych, and Anna Tramontano. Critical assessment of methods of protein structure prediction (casp)—round ix. *Proteins: Structure*, Function, and Bioinformatics, 79(S10):1–5, 2011.

John Moult, Krzysztof Fidelis, Andriy Kryshtafovych, Torsten Schwede, and Anna Tramontano. Critical assessment of methods of protein structure prediction (casp)—round x. *Proteins: Structure*, Function, and Bioinformatics, 82:1–6, 2014.

Lisa D Muiznieks and Fred W Keeley. Molecular assembly and mechanical properties of the extracellular matrix: A fibrous protein perspective. Biochimica et Biophysica Acta (BBA)-Molecular Basis of Disease, 1832 (7):866–875, 2013.

Erik Nijkamp, Jeffrey A Ruffolo, Eli N Weinstein, Nikhil Naik, and Ali Madani. Progen2: exploring the boundaries of protein language models. Cell systems, 14(11):968–978, 2023.

Mansheej Paul, Surya Ganguli, and Gintare Karolina Dziugaite. Deep learning on a data diet: Finding important examples early in training. Advances in neural information processing systems, 34:20596–20607, 2021.

Guilherme Penedo, Quentin Malartic, Daniel Hesslow, Ruxandra Cojocaru, Alessandro Cappelli, Hamza Alobeidli, Baptiste Pannier, Ebtesam Almazrouei, and Julien Launay. The refinedweb dataset for falcon llm: outperforming curated corpora with web data, and web data only. arXiv preprint arXiv:2306.01116, 2023.

Guilherme Penedo, Hynek Kydlíček, Anton Lozhkov, Margaret Mitchell, Colin Raffel, Leandro Von Werra, Thomas Wolf, et al. The fineweb datasets: Decanting the web for the finest text data at scale. arXiv preprint arXiv:2406.17557, 2024.

Felipe Garcia Quiroz, Vincent F Fiore, John Levorse, Lisa Polak, Ellen Wong, H Amalia Pasolli, and Elaine Fuchs. Liquid-liquid phase separation drives skin barrier formation. Science, 367(6483):eaax9554, 2020.

Samyam Rajbhandari, Jeff Rasley, Olatunji Ruwase, and Yuxiong He. Zero: Memory optimizations toward training trillion parameter models. In SC20: International Conference for High Performance Computing, Networking, Storage and Analysis, pages 1–16. IEEE, 2020.

Roshan Rao, Joshua Meier, Tom Sercu, Sergey Ovchinnikov, and Alexander Rives. Transformer protein language models are unsupervised structure learners. bioRxiv, 2020. doi: 10.1101/2020.12.15.422761. URL https://www.biorxiv.org/content/early/2020/12/15/2020.12.15.422761.

Alexander Rives, Joshua Meier, Tom Sercu, Siddharth Goyal, Zeming Lin, Jason Liu, Demi Guo, Myle Ott, C Lawrence Zitnick, Jerry Ma, et al. Biological structure and function emerge from scaling unsupervised learning to 250 million protein sequences. Proceedings of the National Academy of Sciences, 118(15): e2016239118, 2021.

Kiersten M Ruff and Rohit V Pappu. Alphafold and implications for intrinsically disordered proteins. Journal of molecular biology, 433(20):167208, 2021.

Benjamin R Sabari, Alessandra Dall’Agnese, Ann Boija, Isaac A Klein, Eliot L Coffey, Krishna Shrinivas, Brian J Abraham, Nancy M Hannett, Alicia V Zamudio, John C Manteiga, et al. Coactivator condensation at super-enhancers links phase separation and gene control. Science, 361(6400):eaar3958, 2018.

Andrew W Senior, Richard Evans, John Jumper, James Kirkpatrick, Laurent Sifre, Tim Green, Chongli Qin, Augustin Žídek, Alexander WR Nelson, Alex Bridgland, et al. Improved protein structure prediction using potentials from deep learning. Nature, 577(7792):706–710, 2020.

Mathieu Seppey, Mosè Manni, and Evgeny M Zdobnov. Busco: assessing genome assembly and annotation completeness. Gene prediction: methods and protocols, pages 227–245, 2019.

Noam Shazeer. Glu variants improve transformer. arXiv preprint arXiv:2002.05202, 2020.

Mary Shelley. Frankenstein: the 1818 text. Penguin, 2018.

Yongdae Shin and Clifford P Brangwynne. Liquid phase condensation in cell physiology and disease. Science, 357(6357):eaaf4382, 2017.

Yongdae Shin, Yi-Che Chang, Daniel SW Lee, Joel Berry, David W Sanders, Pierre Ronceray, Ned S Wingreen, Mikko Haataja, and Clifford P Brangwynne. Liquid nuclear condensates mechanically sense and restructure the genome. Cell, 175(6):1481–1491, 2018.

A. Shrake and J.A. Rupley. Environment and exposure to solvent of protein atoms. lysozyme and insulin. Journal of Molecular Biology, 79(2):351–371, 1973. ISSN 0022-2836. doi: 10.1016/0022-2836(73)90011-9. URL https://www.sciencedirect.com/science/article/pii/0022283673900119.

Felipe A Simão, Robert M Waterhouse, Panagiotis Ioannidis, Evgenia V Kriventseva, and Evgeny M Zdobnov. Busco: assessing genome assembly and annotation completeness with single-copy orthologs. Bioinformatics, 31(19):3210–3212, 2015.

Ben Sorscher, Robert Geirhos, Shashank Shekhar, Surya Ganguli, and Ari Morcos. Beyond neural scaling laws: beating power law scaling via data pruning. Advances in Neural Information Processing Systems, 35: 19523–19536, 2022.

Martin Steinegger and Johannes Söding. Mmseqs2 enables sensitive protein sequence searching for the analysis of massive data sets. Nature biotechnology, 35(11):1026–1028, 2017.

Boris Steipe, Britta Schiller, Andreas Plückthun, and Stefan Steinbacher. Sequence statistics reliably predict stabilizing mutations in a protein domain, 1994.

Jianlin Su, Murtadha Ahmed, Yu Lu, Shengfeng Pan, Wen Bo, and Yunfeng Liu. Roformer: Enhanced transformer with rotary position embedding. Neurocomputing, 568:127063, 2024. ISSN 0925-2312. doi: 10.1016/j.neucom.2023.127063. URL https://www.sciencedirect.com/science/article/pii/S0925231223011864.

Palaniraja Thandapani, Timothy R O’Connor, Timothy L Bailey, and Stéphane Richard. Defining the rgg/rg motif. Molecular cell, 50(5):613–623, 2013.

Hugo Touvron, Louis Martin, Kevin Stone, Peter Albert, Amjad Almahairi, Yasmine Babaei, Nikolay Bashlykov, Soumya Batra, Prajjwal Bhargava, Shruti Bhosale, et al. Llama 2: Open foundation and fine-tuned chat models. arXiv preprint arXiv:2307.09288, 2023.

Wouter G. Touw, Coos Baakman, Jon Black, Tim A. H. te Beek, E. Krieger, Robbie P. Joosten, and Gert Vriend. A series of pdb-related databanks for everyday needs. Nucleic Acids Research, 43(D1):D364–D368, 10 2014. ISSN 0305-1048. doi: 10.1093/nar/gku1028.

Jesse Vig, Ali Madani, Lav R Varshney, Caiming Xiong, Richard Socher, and Nazneen Fatema Rajani. Bertology meets biology: Interpreting attention in protein language models. arXiv preprint arXiv:2006.15222, 2020.

Jue Wang, Sidney Lisanza, David Juergens, Doug Tischer, Joseph L Watson, Karla M Castro, Robert Ragotte, Amijai Saragovi, Lukas F Milles, Minkyung Baek, et al. Scaffolding protein functional sites using deep learning. Science, 377(6604):387–394, 2022.

Ray Yu-Ruei Wang, Yan Han, Kristina Krassovsky, William Sheffler, Michael Tyka, and David Baker. Modeling disordered regions in proteins using rosetta. PloS one, 6(7):e22060, 2011.

Joseph L Watson, David Juergens, Nathaniel R Bennett, Brian L Trippe, Jason Yim, Helen E Eisenach, Woody Ahern, Andrew J Borst, Robert J Ragotte, Lukas F Milles, et al. De novo design of protein structure and function with rfdiffusion. Nature, 620(7976):1089–1100, 2023.

Bruce J Wittmann, Kadina E Johnston, Zachary Wu, and Frances H Arnold. Advances in machine learning for directed evolution. Current opinion in structural biology, 69:11–18, 2021.

Qian Xiao, Ceara K McAtee, and Xiaolei Su. Phase separation in immune signalling. Nature Reviews Immunology, 22(3):188–199, 2022.

Ruibin Xiong, Yunchang Yang, Di He, Kai Zheng, Shuxin Zheng, Chen Xing, Huishuai Zhang, Yanyan Lan, Liwei Wang, and Tie-Yan Liu. On layer normalization in the transformer architecture. In Proceedings of the 37th International Conference on Machine Learning, ICML’20. JMLR.org, 2020.

Biao Zhang and Rico Sennrich. Root mean square layer normalization. Curran Associates Inc., Red Hook, NY, USA, 2019.

Zhidian Zhang, Hannah K. Wayment-Steele, Garyk Brixi, Haobo Wang, Dorothee Kern, and Sergey Ovchinnikov. Protein language models learn evolutionary statistics of interacting sequence motifs. Proceedings of the National Academy of Sciences, 121(45):e2406285121, 2024. doi: 10.1073/pnas.2406285121. URL https://www.pnas.org/doi/abs/10.1073/pnas.2406285121.

